# The African mosquito-borne diseasosome: Geographical patterns and range expansion

**DOI:** 10.1101/2021.12.21.473756

**Authors:** Tovi Lehmann, Cedric Kouam, Joshua Woo, Mawlouth Diallo, Richard Wilkerson, Yvonne-Marie Linton

## Abstract

Mosquito-borne diseases (MBDs) such as malaria, dengue, and Rift Valley fever threaten public health and food security globally. Despite their cohesive nature, they are typically treated as distinct entities. Applying biological system analysis to the African MBDs from a One Health perspective, we provide the first biogeographic description of the African mosquito fauna corresponding with the pathogens they transmit. After compiling records accumulated over a century, we find that there are 677 mosquito species in Africa, representing 16 genera, and 151 mosquito-borne pathogens (MBPs) circulating primarily among wild tetrapods, dominated by viruses (95) and protozoans (47). We estimate that reported MBPs represent ∼1% of the actual number. Unlike mosquitoes, African arboviruses and mammalian plasmodia represent a higher share of the World’s total based on the area – species richness relationship (P<0.0001), explaining the disproportional large share of global MBPs that originated from Africa. Species richness of African mosquitoes and MBPs are similarly concentrated along the equator, peaking in central Africa, with a secondary “ridge” along eastern Africa. Moderate diversity and low endemicity in mosquitoes across the Sahel reveals a fauna with high propensity for long-range migration. Regional differences in species richness, endemicity, and composition agreed with country-based results. The composition of mosquitoes and MBPs separates sub-Saharan Africa from north Africa, in accordance with the Palearctic and Afrotropical faunal realms, and west and central Africa are clustered together distinctly from the cluster of eastern and southern Africa. With ∼25% of the species occupying a single country, ∼50% in 1–3 countries and <5% found in >25 countries, the typical ranges of both mosquitoes and MBPs are surprisingly small. The striking similarity in diversity and especially in range distributions of mosquitoes and MBPs suggest that most MBPs are transmitted by one or few narrow-range mosquito vectors. Exceptionally widespread mosquito species (e.g., *Ae. aegypti, Cx. quinquefasciatus*, and 10 *Anopheles* species) feed preferentially on people and domestic animals, and nearly half are windborne migrants. Likewise, exceptionally widespread MBPs are transmitted between people or domestic animals and are vectored by one or more of the aforementioned widespread mosquitoes. Our results suggest that few MBPs have undergone a dramatic range expansion, after adapting to people or domestic animals as well as to exceptionally-widespread mosquitoes. During the intermediate phase of range expansion, MBPs extend their vector and vertebrate host ranges with a concomitant gradual increase in geographical range. Because range size may serve as a marker of the phase of range expansion, ranking the African MBPs according to range, we identified several MBPs that pose elevated risk for disease emergence (e.g., Wesselsbron virus). Taken together, our database, approach, and results can help improve MBD surveillance and lead to a better understanding of disease emergence. This knowledge has the potential to improve capacity to prevent and mitigate new and emerging MBD threats.

## Introduction

Africa carries the heaviest global burden of mosquito borne diseases (MBDs), with more than 400,000 deaths attributable to malaria of the 700,000 caused by vector-borne diseases annually (WHO, 2020). At least eight of the 11 topmost impactful global mosquito-borne pathogens (MBPs) originated in Africa—namely Yellow Fever virus (YFV), West Nile virus (WNV), Chikungunya virus (CHIKV), Rift Valley fever virus (RVFV), Zika virus (ZIKV) and three human *Plasmodium* species (*falciparum, malariae, ovale*) (WHO, 2020). Excluding its islands, Africa comprises only 20% of the earth’s land surface, but is the origin of 73% (8 of 11) of these global MBPs. Based on the species richness – area relationship (Lomolino, 2020), this excess is highly significant (P<0.001, exact binomial test), corroborating a recent literature review that has reached a similar conclusion using different data (Swei *et al*., 2020). The reasons for Africa’s disproportional role as the origin of so many global MBPs may include being the only continent that extends from the northern to southern temperate zones, covering >70° of latitudes (37°N– 34°S), and straddling several biomes including the outstandingly diverse equatorial forest (Burgess *et al*., 2004; Guernier *et al*., 2004; Lomolino, 2020). As the homeland of the hominids and extant apes, we expected that Africa would contain most human MBPs (Wolfe *et al*., 2007), yet, five of the eight global MBDs are zoonotic (YF, WN, RVF, CHIK, and ZIK), leading to a new hypothesis that Africa has more MBDs in total, not only those affecting humans. Africa is also home to the largest number of megafauna species, and thus it poses a greater risk to many phylogenetically related domestic animals. Understanding the MBDs of Africa, therefore, would be valuable for global health and food security. As Africa undergoes dramatic perturbations due to deforestation and global climate change (e.g., desertification), coupled with food and water scarcity, the risk for the emergence/re-emergence of MBDs should be closely monitored. Baseline knowledge is a prerequisite to successful mitigation of MBDs, however– as we have found–this vital information is scarce and not readily accessible.

The study of MBDs has traditionally been fragmented into separate fields: virology, parasitology, entomology, etc. Most studies have focused on one or a few related pathogens, and/or their vectors in a limited region. Excepting a few reviews of certain MBDs (Weaver *et al*., 2012; Braack *et al*., 2018), the ensemble of MBDs as a biological system composed of mosquitoes and pathogens has never before been holistically studied to our knowledge. On the other hand, the increasing frequency of disease emergence in humans has deservedly been the focus of extensive study (Burke, 1998; Binder *et al*., 1999; Taylor *et al*., 2001; Woolhouse & Gowtage-Sequeria, 2005; Jones *et al*., 2008; Morse *et al*., 2012; Rosenberg, 2015), yet their broad scope may have precluded inferences into commonalities shared among certain groups of diseases. Here, focusing on the MBDs in continental Africa from a One Health perspective, we include all known pathogens transmitted by mosquitoes to terrestrial tetrapods, and compare biogeographical features of this disease system after constructing a dedicated database, based on a comprehensive literature search (Supp. Materials). We describe the composition and geographical organization of the mosquito species and the MBPs in Africa to better understand the process of MBD range expansion. Specifically, we evaluate the hypothesis that Africa has exceptionally high mosquito and MBP diversities, and map the landscapes of their species richness, endemicity, and composition. The results lead to insights into the role of mosquito and MBP dispersal, the nature of barriers to their spread, and the future of MBD surveillance in Africa. Evaluating variation in range size of mosquito and MBP species and attributes of extremely widespread species, we propose a process for range expansion of MBDs and accordingly rank the African MBDs as to their present phase of range expansion as a marker of risk for disease emergence.

Despite being studied for over a century, the main vector species of MBPs of vertebrates remain largely unknown, including many sylvatic vectors (transmitting among wild animals) of the most well-studied pathogens (Karabatsos, 1985; Service, 2001; Njabo *et al*., 2009; Diallo *et al*., 2012; Perkins, 2014; Kyalo *et al*., 2017; Villinger *et al*., 2017; Nanfack Minkeu & Vernick, 2018; Weaver *et al*., 2020; Wilkerson *et al*., 2021). This is also the case for many MBP species of vertebrates (Karabatsos, 1985; Njabo *et al*., 2009; Weaver *et al*., 2012; Perkins, 2014, 2018). Therefore, it is most likely that the role of many mosquito species as vectors of known and unknown pathogens is yet to be discovered. Accordingly, we included all known African mosquitoes as the basis for describing and understanding patterns that we expect would also apply to as yet unrecognized mosquito vector species. In this exploratory analysis, we summarize trends based on knowledge that has been accumulated over at least 120 years. With the expected growth in this domain due to the renewed recognition of the value of disease surveillance brought about by the ongoing COVID-19 pandemic, as well as the advance in methodologies such as metagenomics, it would be valuable to revisit these trends every decade and assess the changes in the patterns described herein.

## RESULTS

As many of the records on mosquito and MBP distribution were collected before 1980, reliable localization of a large portion of these records is only available at the country level (Karabatsos, 1985; Fontenille *et al*., 1998; Foley *et al*., 2007; Kyalo *et al*., 2017; Braack *et al*., 2018; CDC, 2019; Irish *et al*., 2020; Wilkerson *et al*., 2021). Because many African countries cover multiple ecozones (Burgess *et al*., 2004) and biogeographic regions as defined for various animal classes (Linder *et al*., 2012), and vary in size, our analysis addresses fuzzy eco-geographic units.

### Composition of African mosquitoes and mosquito-borne pathogens

Continental Africa, which comprises 20% of the Worlds land surface, supports 19% of all known global mosquito species (N = 3,570, see Methods) (Wilkerson *et al*., 2021). The African mosquito fauna is represented by 677 species spanning 16 genera and 53 subgenera, with *Aedes* comprising the largest number of species, followed by *Anopheles* and *Culex* (Fig. 1a). Goodness of fit tests of 20% across the 16 represented genera where N >15 species/genus revealed higher fractions of African species in *Aedes* (23%, P<0.01, X^2^_1_=6.9), in *Anopheles* (30%, P<0.0001, X^2^_1_ = 27.5), and in *Coquillettidia* (40%, P<0.0002, X^2^_1_=14.0), but was insignificant in the other genera. The highest fraction of African species (100%) is found in the genus *Eretmapodites* (n = 48), which is endemic to Africa (Fig. 1a). Fractions >20% were found in several genera, indicating local speciation on the continent. Among the 53 mosquito subgenera in Africa, *Anopheles* subgenus *Cellia* is by far the most speciose (n = 121, Fig. 1b, Table S1). Several subgenera have a high proportion of African species (Fig. 1b), although most of these have a small number of species in total, e.g., *Anopheles* subgenus *Christya* (n= 2). Nonetheless, the 29 *Aedes* species in the subgenus *Catageiomyia* are exclusively African species and 24 of the 28 species of the *Aedes* subgenus *Neomelaniconion* are African (Fig. 1b, Table S1). While not precluding that some of the species also occur outside Africa, a high fraction of species found in Africa, especially in taxa with large number of species, highlights the fauna’s unique elements.

**Fig. 1:**
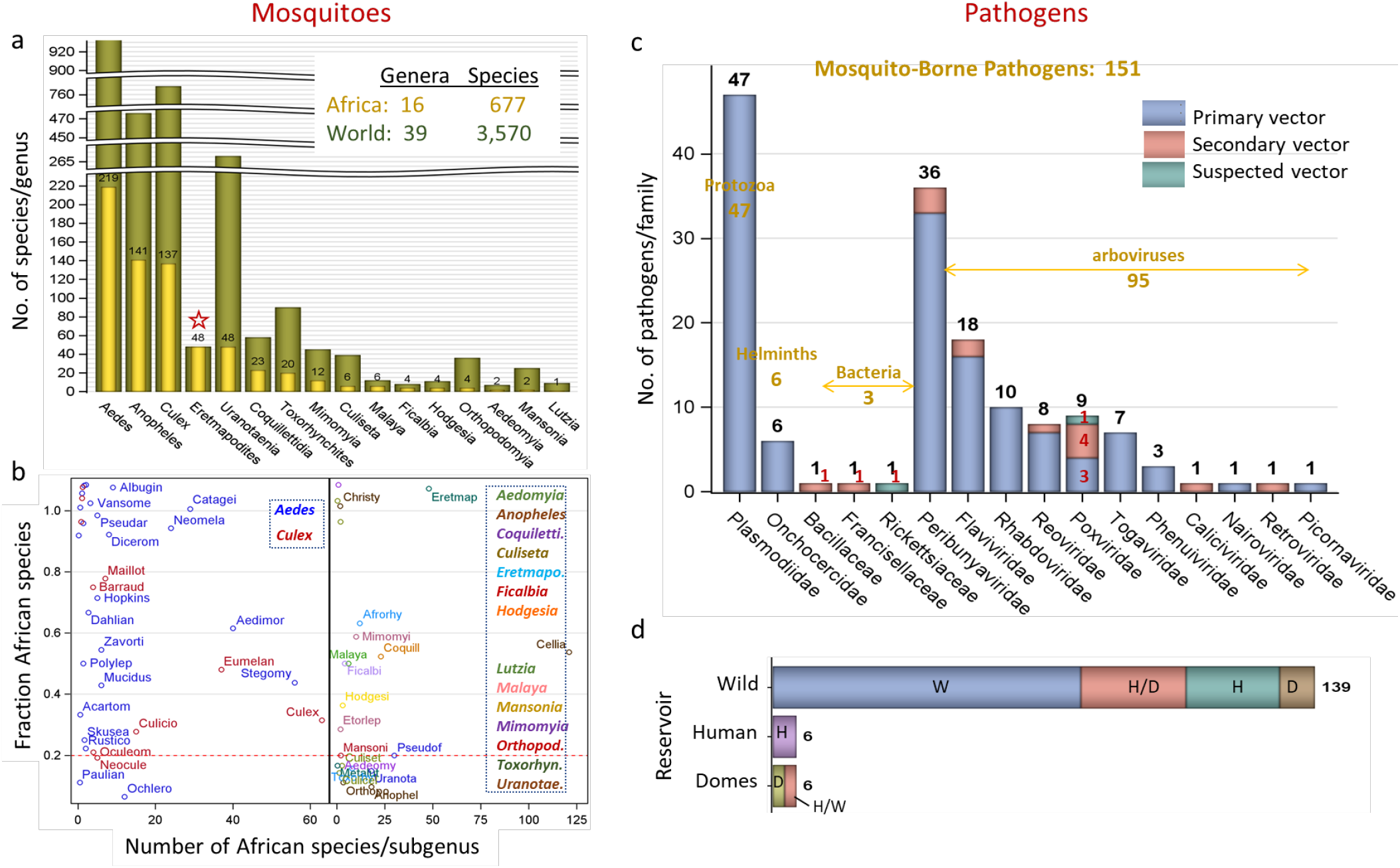
Taxonomical composition of African mosquitoes and MBPs. a) Number of African species/genus (gold and numerals) compared to the total number worldwide (green). Star denotes entirely African genus. Note: breaks in the Y-axis. b) The fraction of African mosquito species per subgenus (Y-axis) of their worldwide total in relation to their number in Africa (X-axis). To minimize label overlap, values near 1 were jittered. Subgenera labels (abbreviated to the first 7 letters) are shown if they have two or more species. Where no subgenera are known, e.g., *Ficalbia* (Table S1), genus names were used. Corresponding genera (bold italics font) of the same color are listed in dotted frame. Red line marks expected 20% based on African share of land surface (text and Suppl. Table 1). c) Taxonomic composition of African mosquito borne pathogens affecting vertebrates by family and importance of mosquito-borne transmission (legend). Suspected mosquito transmission reflects compelling, yet non-definitive evidence (Supp. Mat). The number of pathogens in each family is shown above bars (black) and the total by taxonomic group shown across (gold). The number of pathogens transmitted mechanically are listed (red). d) Division of MBPs by group of vertebrate host acting as reservoir (Y-axis) and by the host group impacted by the pathogen (subgroups in color). Key: W, H, and D denote wild, human, and domestic animal (those raised by people, Domes), respectively and H/D denotes that humans and domestic animals are impacted by MBPs whose reservoir are wild animals.

A total of 151 known mosquito-borne-pathogen species (MBPs) affecting vertebrates have been reported from continental Africa (Fig. 1c). These include 95 viruses, 47 protozoans, six helminths, and three bacteria, comprising a total of 16 families and 30 genera (Fig. 1c, including 3 unclassified genera). These 95 arboviruses represent a significantly higher share than expected from the known global total based on the surface land area of Africa (32% vs. 20%, P<0.0001, X^2^_1_=25.5). The fraction of MB arboviruses is likely even higher because among the total of 300 arboviruses that have been isolated from mosquito pools worldwide (Wilkerson *et al*., 2021), some are probably not vectored by mosquitoes. Likewise, of the 60 mammalian plasmodia (Perkins 2014), 27 species (40%) are reported from Africa, which is a larger than expected based on the continental/global land mass area (P<0.0001, X^2^_1_=23.4). Plasmodia of birds were not considered here because recent molecular analyses revealed many lineages that likely represent yet-to-be named species (Bensch *et al*., 2009; Njabo *et al*., 2009; Perkins, 2014). Additionally, many trans-continental migrant birds may be exposed to plasmodia only in Europe and Asia (Hellgren *et al*., 2007) and thus should not be considered African — a problem that may apply to other avian pathogens.

The vast majority of MBPs are maintained in wild hosts reservoirs (Fig. 1d), although a few can be transmitted for short periods between humans, e.g., YFV, O’nyong’nyong virus (ONNV) or domestic animals e.g., RVFV. Mosquito transmission is the primary route of vertebrate infection in the majority of the MBPs (Fig. 1c), whereas 16 pathogens rely on other arthropods or on direct transmission as their primary mode of transmission and use mosquitoes as a secondary route (Fig 1c). At least 8 of the 9 poxviruses are transmitted mechanically by mosquitoes as are the three bacteria (Turell & Knudson, 1987) (Fig.1c, Supp. File 2). Mechanical transmission appears to be linked to secondary transmission (Fig. 1c, Supp. File 2), although certain poxviruses can be transmitted several weeks post exposure (Kligler *et al*., 1928; DaMassa, 1966).

#### Range size occupied by the African mosquitoes and MBPs

Crude range approximations defined by the number of African countries occupied by mosquito species revealed that 26% of all African mosquito species are endemic (known only within a single country), and that over 50% are restricted to only 1–3 countries (median= 3.0, Fig. 2a). The L-shape distribution reveals that only 5% of the total number of species are widespread across over half the continent (i.e., occupying 26 or more countries, Fig. 2a). The Pearson correlation coefficient between the number of countries and total area (sum over countries areas) occupied by each species is 0.966 (N= 677, P<0.0001), indicating that the number of countries approximates range size reasonably well. The median area occupied by a mosquito species is 3.09 ×10^6^ km^2^ (95% CI: 2.72–3.42 km^2^). Range size varies among genera (P<0.05, quantile regression, Fig. 2a: inset).

**Figure 2.**
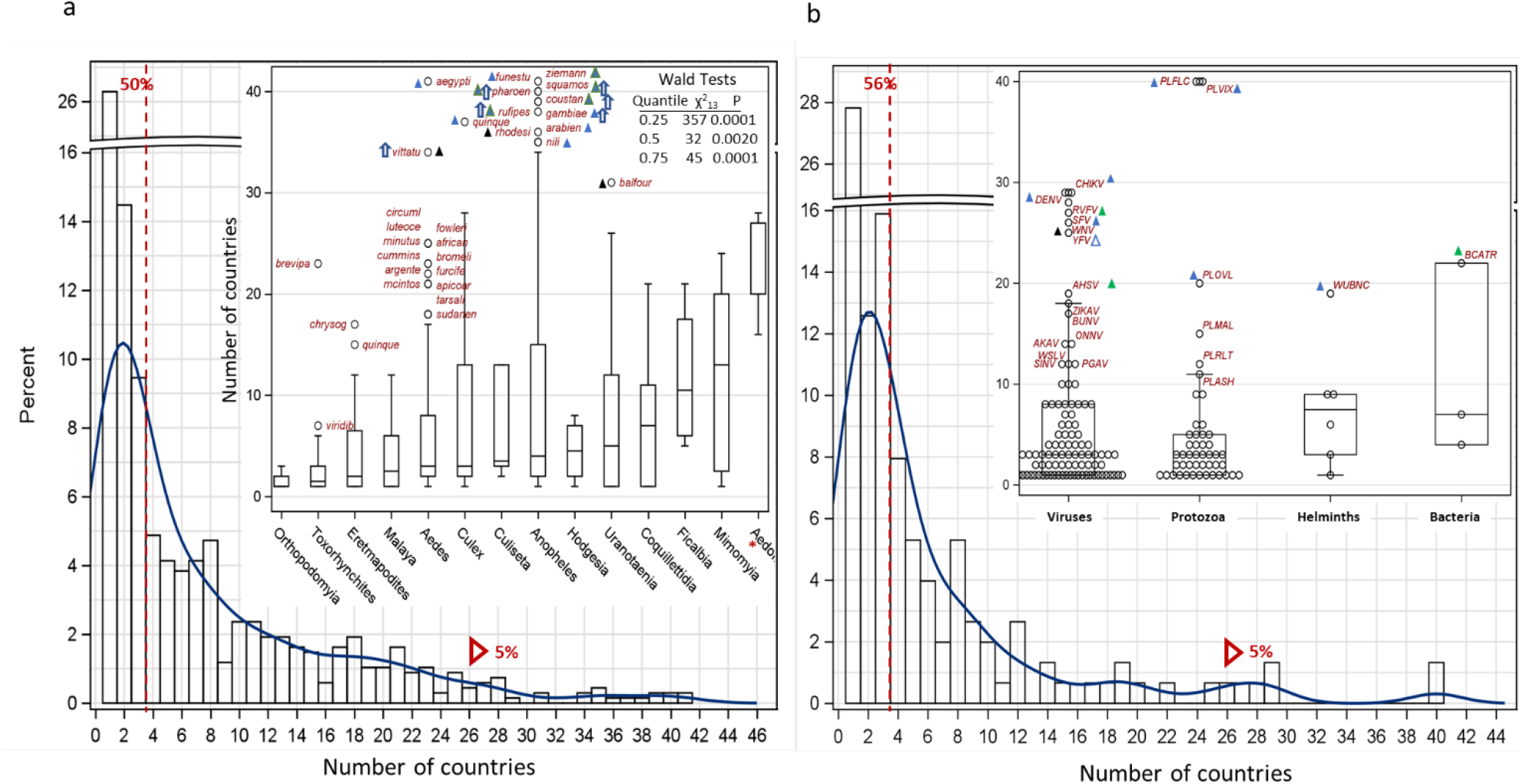
Mosquito (a) and MBPs (b) geographic range based on the number of countries per species overall and by taxonomical groups (Insets). Note the break in the Y-axis. The fraction of species occupying 1–3 countries and over 25 countries are shown to the left of the red broken line and red triangle, respectively. Insets: Number of countries per species across mosquito genera (a) and taxonomic group of MBPs (b). Genera represented by <3 species (*Lutzia* (N=1), *Mansonia* (N=2), and *Aedeomyia* (N=2)) were pooled (red asterisk). The box shows the 25^th^, 50^th^, and 75^th^ quantiles of the distribution and the whiskers extends to the more extreme observations up to 1.5 x the interquartile range (75^th^ – 25^th^ quantiles). Outliers exceeding the whiskers are shown by abbreviated species name (a) and acronym (b) in red; triangles indicate preference to blood feeding on (a) and transmission between (b) humans (blue) domestic animals (green) and wild hosts (black). Empty triangle (b) indicate transmission to humans but no persistent transmission from humans. Table (a) summarizes results of the quantile regression (see text).

Species of *Lutzia, Mansonia, Aedeomyia, Mimomiya, Ficalbia*, and *Coquillettidia* have the largest ranges (Fig. 2a). However, the most widespread mosquito species that are found in 30 or more countries (14 of 677 species, Fig. 2) include *An. gambiae, An. arabiensis, An. funestus, Ae. aegypti* and *Cx. quinquefasciatus*, as well as *An. pharoensis, An. squamosus, An. coustani, An. ziemanni* and *Ae. vitattus*. With ten of the 14 species, *Anopheles* predominates this group of exceptionally widespread species. Eleven of these 14 species thrive in domestic environments and blood-feed on people or domestic animals (Fig. 2a), whereas at least six engage in high altitudes wind-borne migration based on recent studies in Mali (Huestis *et al*., 2019).

The distribution of African MBPs is remarkably similar to that of the mosquitoes (Fig. 2b), with 28% being single country-endemic, 56% found in 1–3 countries (median= 3.0), and 5% found in 26 or more countries (i.e., over approximately half the continent). The Pearson correlation coefficient between the number of countries and total area occupied by each MBP species is 0.967 (N= 151, P<0.0001), corroborating the suitability of the number of countries as an index of total area. The median area occupied by a MBP is 2.15 ×10^6^ km^2^ (95% CI: 1.64-2.65 ×10^6^ km^2^). The most widespread MBPs (40 countries) are *Pl. falciparum and Pl. vivax* (PLFLC, PLVIX) with only nine MBPs being reported in 20 or more countries (Fig. 2b). Except West Nile virus (WNV), which is primarily transmitted among birds (including migratory birds), all of these exceptionally widespread MBPs are transmitted among humans (6) or domestic animals (2), and without exception, all are vectored by at least one of the most widespread mosquito species mentioned above (Fig. 2a). For example, PLFLC, PLVIX, and *Pl. ovale* (PLOVL) are transmitted by all, or some of the above *Anopheles* species, and DENV, CHIKV, and YFV are transmitted by *Ae. aegypti* in urban and semi-urban settings (Jupp & McIntosh, 1990; Collins & Jeffery, 2005; Diallo *et al*., 2014; Faye *et al*., 2014; Kyalo *et al*., 2017; Braack *et al*., 2018; Twohig *et al*., 2019). Similarly, WNV is transmitted by *Cx. quinquefasciatus* and YFV and ZIKV are transmitted by *Ae. vittatus* (Faulde *et al*., 2012; Epelboin *et al*., 2017; Braack *et al*., 2018; Diagne *et al*., 2021).

#### Diversity and endemism across the continent

According to the area - species richness principle (Lomolino, 2020), mosquito species richness has been found to increase with a country’s area in a worldwide analysis (Foley *et al*., 2007). In Africa, this relationship accounted for only 13% of the variance compared to 42% worldwide, highlighting the importance of other factors (Fig. 3a). The regions of highest species richness include a belt along the equatorial forest, which appears widest in central Africa (Fig. 3a). The countries with highest species richness include DRC, Cameroon, Uganda, Kenya, Nigeria, and Ivory Coast. Adjusting for area minimally changes these countries’ ranking (Fig. 3a). North Africa represents a uniform belt of lowest mosquito diversity (Figs. 3a) with Libya being an extreme outlier that exceeds the 95% CL, given its area (Fig. 3a). A corridor of modest diversity along the Sahel (from Mauritania to Chad), and possibly another corridor between central and East Africa includes countries from Namibia and Botswana to Rwanda, which remains stable after accommodating country area (Fig. 3a).

**Figure 3.**
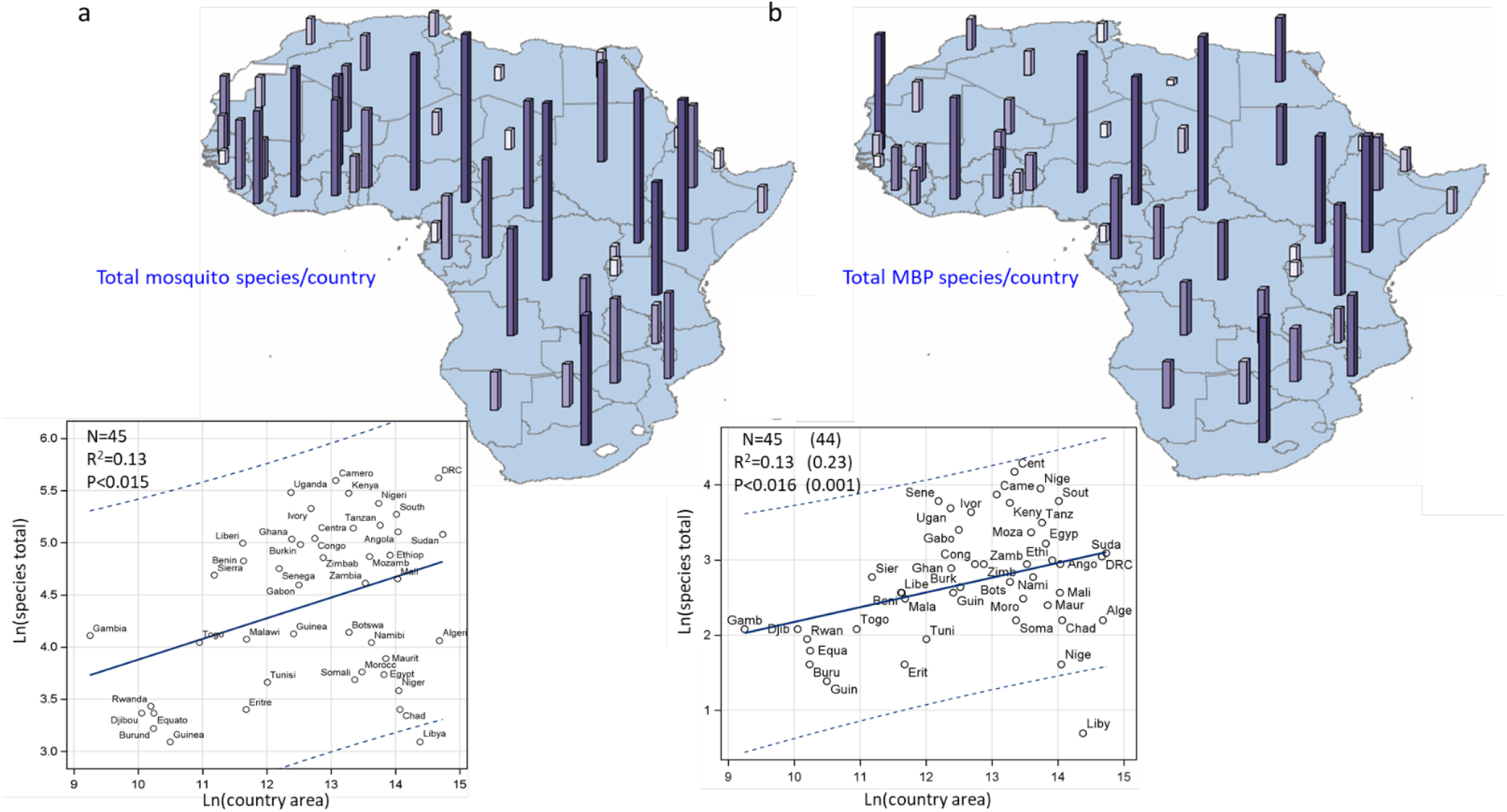
Maps showing mosquito (a) and MBP (b) country diversity (top) and the country’s area – species richness relationship (bottom). Bar height and color shows total endemic species per country (top). Linear regression (solid line) shows the increase in expected number of species per country given its area with 95% confidence limit for individual countries (broken lines). Values in parenthesis (b) show the change in the regression’s summary statistics after exclusion of outliers. Abbreviated country names are used. Note: Countries with insufficient information are excluded (e.g., Eswatini), or pooled together (e.g., South and North Sudan) to reflect available information (Methods).

Similar to mosquitoes, species richness of African MBPs is highest in Central Africa, followed by an East African zone stretching from Kenya to South Africa (Fig. 3b). Except Senegal and the Ivory Coast, West Africa exhibits lower diversity of MBPs than East Africa. Sahelian countries and those between Central and East Africa exhibit lower MBP richness than the surrounding regions, whereas North Africa exhibits the lowest MBP richness (Fig. 3b). Species richness increased with country’s size, but this relationship accounted for only 13% of the variance among countries (excluding outliers: Libya, increased R^2^ to 23%, Fig. 3b).

The distribution of country endemic mosquito species reveals greater heterogeneity than species richness, with highest endemicity in Equatorial Central Africa, especially Cameroon (31), followed by South Africa and Angola (22, Fig. 4a). These three countries represent outliers after accommodating species richness and, indirectly, country area (captured by species richness, Fig. 3a). Unlike species richness, lowest mosquito endemicity is found across the Sahel from Mauritania to Somalia and, notably, extending to equatorial West Africa. Additionally, the secondary “corridor” of low species richness separating South Africa from Central and East Africa (Fig. 3a) appears wider for endemicity. Countries without known endemic mosquito species included Chad and Mozambique (Fig. 4a). The number of endemic species per country is correlated with the total species richness (r= 0.7, N= 45, P< 0.001, Fig. 4a), but the relationship was not monotonic, and visual inspection suggests a higher slope after species richness exceeds ∼100 species per country (Figs. 3a and 4a).

**Figure 4.**
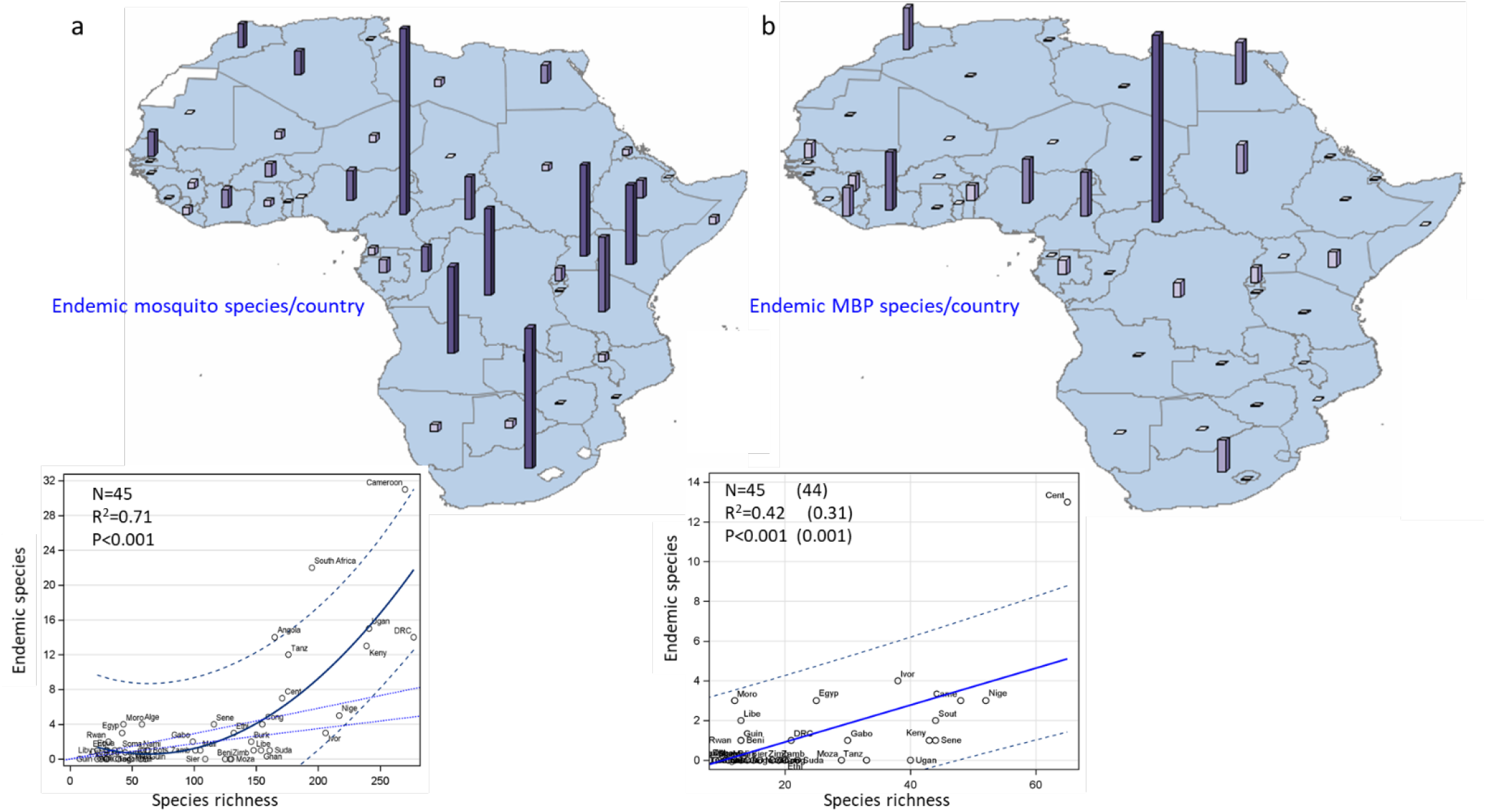
Maps showing endemic species of mosquito (a) and MBP (b) per country (top) and the relationship between species endemicity and richness (bottom). Bar height and color shows total endemic species per country (top). Quadratic (a, bottom) and linear regression (b, bottom) solid lines show the increase in expected mean number of endemic species given species richness with 95% confidence limits for individual countries (broken lines). Dotted lines show expected linear trends assuming monotonic increase predicted by the mean (higher) and median (lower) ratio of endemics to total species. Values in parenthesis (b) show the regression summary statistics after exclusion of outliers. Abbreviated country names are used.

Country-endemic MBPs comprised 25 arboviruses, 11 plasmodia, and 1 nematode, reflecting similar proportion of endemicity across taxa: 27.5%, 25.6%, and 16.7%, respectively. Endemic MBPs showed an extreme hotspot in Central African Republic, moderate endemism in Ivory coast, followed by Nigeria and Cameroon, Egypt, and Morocco (Fig. 4b). Endemic MBP species per country also increased with species richness and indirectly with species area, which was a determinant of the latter (Fig. 3b). After accounting for species richness, the Central African Republic remains an outlier endemic hotspot, towering over all other countries.

### Heterogeneity in species composition across regional and country scales

Because countries differ considerably in surveillance effort, a regional analysis, whereby each region consists of multiple countries, exhibits less variability in surveillance effort and can be used to ascertain the patterns noted at the country level. Five regions were defined to group neighboring countries together and maximize distances among regions, accommodate latitudinal variation, and minimize inter-region enclaves (without regard to political regions, specific ecological, or species distributional data, Methods). Over 40% of the mosquito species are region-endemic and 60% of the mosquito species are found in 1-2 regions. Only 19% are found across sub-Saharan Africa and merely 3% are distributed across the five regions (Fig. 5a: histogram). Consistent with country results (Fig. 4a), the highest mosquito richness and endemicity are found in Central Africa and the lowest species richness is in North Africa. Notably, endemicity is lowest in West Africa (Fig. 5a). Significant excesses of species richness based on the region size were found in West and Central Africa, whereas a significant deficit was found in North Africa (P<0.01, Z> 3.1, Exact Binomial tests, Fig. 6a). Significant excess of endemic species was detected in Central Africa, whereas West Africa exhibits the lowest endemicity and largest deficit of endemic species based on its size, reflecting the large number of species it shares with Central Africa (N= 57), as well as with both East and Central Africa (N= 60, Fig. 5a).

**Figure 5.**
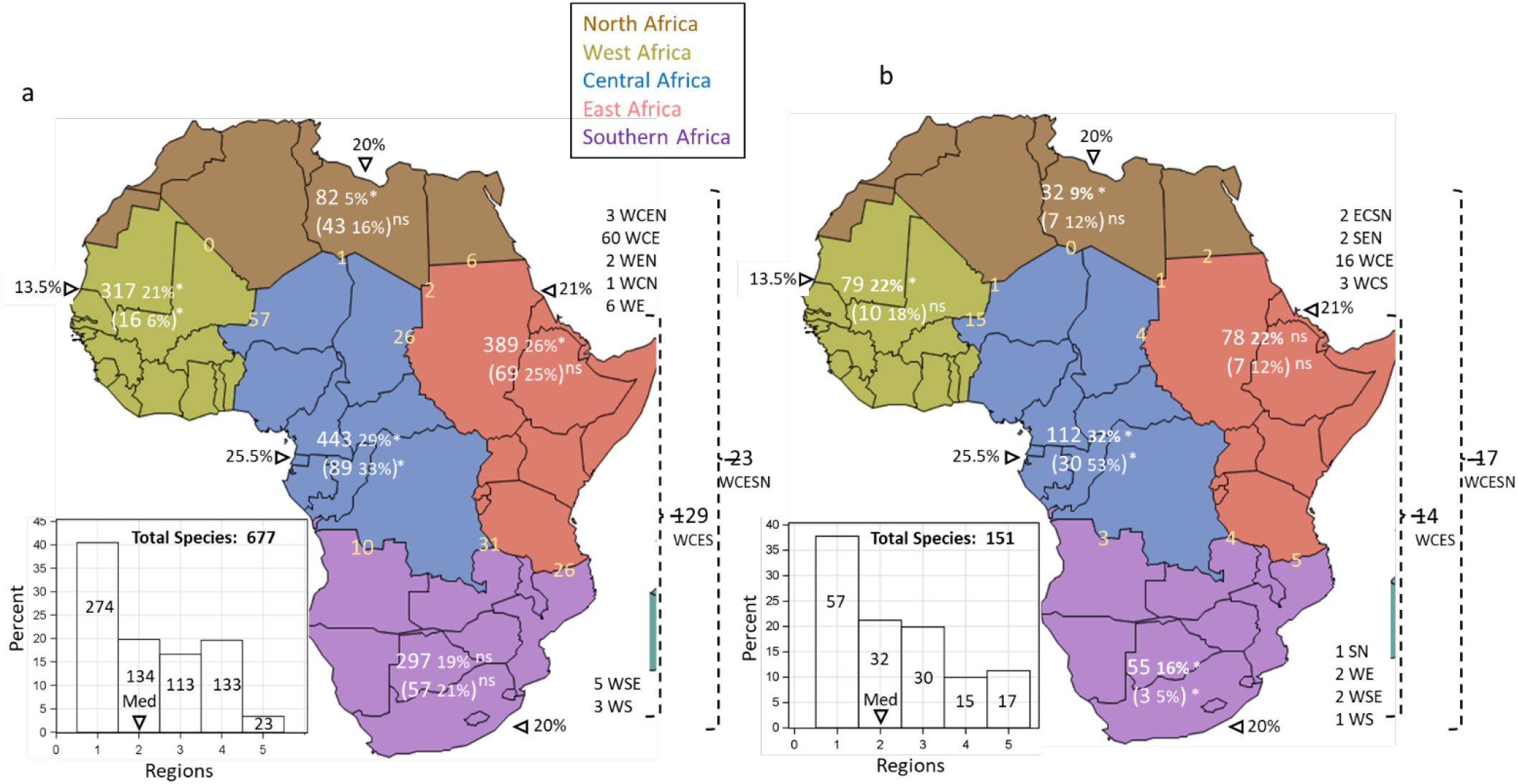
Composition of mosquitoes (a) and MBP species (b) across 5 regions in Africa (North, West, Central, East and Southern Africa). Regional species richness and (region-endemic) in absolute and percentage numbers (white font) against each region’s relative landmass in percent (black font behind triangles). Statistically significant departures (P<0.05) of the actual species richness or endemicity from expectations based on the region area is marked by ‘*’; ‘ns’ denotes insignificant departure. The number of species between two and three adjacent regions are shown at the border between regions (yellow font). Number of species shared between disjoined regions are shown on the right (black) before the regions’ acronym. Histogram showing the number of species occupying different number of regions (median= 2, marked by a black triangle, mean= 2.25).

**Figure 6.**
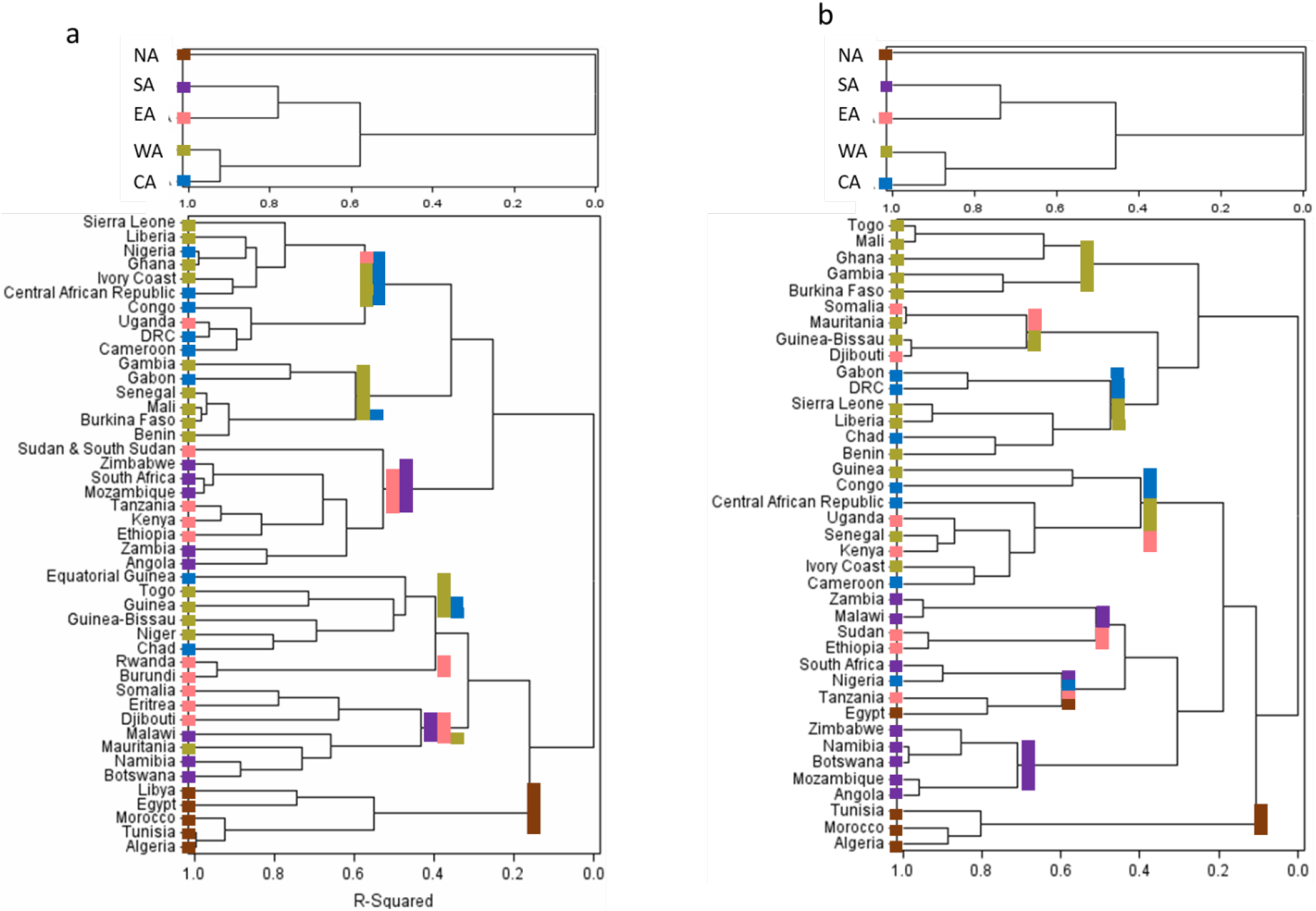
Dendrograms showing clustering of regions (above) and countries (below) based on species composition of mosquitoes (a) and MBPs (b). Region and country color follows color of the regions in Figure 5 (North Africa (brown), West Africa (green), Central Africa (blue), East Africa (pink), Southern Africa (purple)).

Similar to trends seen for mosquitoes, 38% of the MBP species are region-endemic, 60% are found in 1–2 regions, only 10% are found across sub-Saharan Africa, and an additional 11% are found across the continent (Fig. 5b: histogram). MBP richness is highest in Central Africa and lowest in North Africa, whereas West and East Africa have similar MBP richness which appear higher than that of Southern Africa (Fig. 5b). Considering the region’s area, excess of MBPs was detected in Central Africa and West Africa, whereas a deficit was detected in North Africa (Fig. 5b, P<0.01, |Z|> 2.6, Binomial test). MBP endemicity is also highest in Central Africa, but lowest in Southern Africa, showing corresponding sharp departures from expectations based on the region’s area (Fig. 5b, P<0.01, |Z|> 2.5, Binomial test). Similar numbers of region-endemic MBPs are found in West, North, and East Africa in accord with expectations based on area (Fig. 5b, P>0.05). West and Central Africa share more MBPs than other region pairs, whereas North Africa shares the fewest MBPs with all adjoining regions (Fig. 5b). Overlapping MBPs between three regions was especially high between West, Central, and East Africa (N= 16) compared to other combinations (1–4, Fig. 5).

The regional mosquito fauna is split into sub-Sahara and North Africa – the most distinct divisions in term of mosquito species composition, followed by further split of sub-Saharan Africa between the West - Central and the East - Southern fauna (Fig. 6a: top). A country-based dendrogram reveals a more complex picture (Fig. 6a: bottom). Most countries from West and Central Africa are grouped together, as are countries from East and Southern Africa (Fig. 6a). Nine of the twelve high similarity clusters (R^2^>0.9) group countries from the same region, with only three exceptions (Nigeria–Ghana, Ivory Coast–Central African Republic, and DRC– Uganda, Fig. 6a), which share ecological similarity if not geographic continuity. The country dendrogram suggests substantive differences in mosquito faunas between Sahelian and equatorial West Africa countries, which is further supported by the grouping of Chad with Niger, as well as Nigeria with Liberia. Assemblages of mosquitoes defined by their significant co-occurrence in particular areas, independently from the regions defined above are illustrated in Fig. S1a (Supplementary Results and Discussion).

The composition of the MBPs at the regional scale follows almost exactly that of the mosquitoes, showing a deep split into sub-Sahara and North African fauna, followed by a further split of sub-Saharan Africa into the West-Central Africa and the East-Southern African fauna (Fig. 6b, top). The MBP composition of West and Central Africa are most similar. The country-based dendrogram based on MBP composition reveals more pervasive cross-regional and cross sub-divisions clusters. For example, North African countries are clustered together, but Egypt is clustered with Tanzania (Fig. 6b). Nonetheless, most countries are grouped by their region or sub-division, i.e., West and Central African countries as well as East and Southern African countries (Fig. 6b). Departures from regional clustering often follows ecological similarity between countries in the equatorial forests, (e.g., Cameroon and Ivory Coast). Assemblages of MBPs defined by their significant co-occurrence in particular areas, independently from the regions defined above are illustrated in Fig. S1b (Supp. Results and Discussion).

## DISCUSSION

Africa is undergoing explosive growth in human population density, deforestation, desertification, and urbanization – processes that are projected to significantly impact disease dynamics as they increase exposure of humans and domestic animals to diseases of wildlife and generate conditions favoring rapid disease spread (Taylor *et al*., 2001; Guernier *et al*., 2004; Jones *et al*., 2008; Fenollar & Mediannikov, 2018). Given Africa’s high burden of MBDs and the disproportionally high number of global MBDs that originated from the continent, a better knowledge of the African mosquito borne diseasosome, as a biological system, is essential to improve local and global health and food security. To our knowledge, this study provides the first holistic description of the biodiversity of mosquitoes and MBPs in any continent. Despite scarce/incomplete information and historical unbalanced sampling efforts of these taxa across Africa and the globe (below), the data recovered herein summarizes more than a century of bio-surveillance efforts and is worthy of compilation and exploration to guide future surveillance efforts by recognizing key knowledge gaps. Our results identify regions expected to contain more sylvatic vectors of known and yet-to-be discovered MBPs and advance understanding of the factors that have shaped the diversity of the African MBDs. Here, we examine the process of range expansion of these diseases, as a key element of disease emergence and interpret our results by addressing the following key questions: (i) Does exceptional biodiversity of mosquitoes and MBPs in Africa account for its disproportionally larger share as the origin of global MBDs and how likely is this trend to continue? (ii) What is the geographical organization of mosquitoes and MBPs in Africa and has the former structured the latter? and, (iii) What are the roles of domestication, dispersal, and adaptation to new vectors and hosts as drivers of MBD range expansion?

A caveat of our analysis is the low resolution of the country-based distributional data. As explained above, a substantial part of the records on mosquito and MBP distribution is only available at the country level. Because many African countries cover multiple ecozones (Burgess *et al*., 2004) and biogeographic regions (Linder *et al*., 2012), our geographical analysis is coarse, addressing fuzzy geographic-ecological units. Although well-defined geographic-ecological units based on new high-resolution data will improve future biogeographical investigation of the African mosquito-borne diseasosome, our analysis and interpretation of the results accommodate these limitations.

### Global MBDs that originated from Africa: past and future

The disproportional larger share of global MBDs originating from Africa (Swei *et al*., 2020) along with the zoonotic nature of most of these diseases (Introduction), led to our hypothesis that the African mosquito and/or MBP faunas are especially diverse. Our results reveal that the share of the African mosquito fauna (677 species) in the global culicid diversity closely agrees with expectation based on the continental land area (19% vs. 20%). Moreover, it is dominated by cosmopolitan genera, such as *Aedes, Culex*, and *Anopheles* (Fig. 1a), indicating no markedly distinct fauna based on genera composition; yet, it has a distinct assemblage of subgenera (Fig. 1b). Unlike mosquito diversity, the biodiversity of African arboviruses and (mammalian) plasmodia–the largest taxonomic groups of MBPs (Fig. 1c) –are considerably greater than expected by land mass at 30% and 40%, respectively (P<0.0001). These data support a disproportionally higher diversity of MBPs in Africa than in any other continent. Avian plasmodia could not be evaluated as explained above. The higher diversity of African MBPs (but not of African mosquitoes) may account for the higher share of global MBDs originating in Africa and in part for its disproportionally heavy burden of MBDs. While, we cannot rule out the possible effect of greater sampling effort of MBPs in Africa compared with other continents. Yet, a recent study reveals that sampling effort of emergent vector borne diseases has been lower in Africa compared with other continents (Swei *et al*., 2020). If the excess diversity of MBPs reported here is correct, new global MBPs will continue to emerge from Africa at a higher rate than from other continents, making Africa a prime target for future disease surveillance and control.

Unlike the mosquito fauna, which is mostly well described, the African MBPs remain poorly known, given that 47% of the MBPs have been found in humans and domestic animals whereas, at least 92% are maintained in wild species reservoirs (Fig. 1c). A conservative estimate of the African MBPs of vertebrates can be derived assuming that humans represent a typical host for vertebrate-specific MBPs of African origin. As the most thoroughly studied vertebrate, humans are known as the only natural host for at least three and possibly five African plasmodia (Liu *et al*., 2010; Rutledge *et al*., 2017; Arisue *et al*., 2021; Daron *et al*., 2021), and possibly one nematode (Laurence, 1989; Small *et al*., 2019). This may be an underestimate since humans are among the youngest vertebrate species. With over 5,000 vertebrate species in Africa (∼1,400 mammals (Burgin *et al*., 2018), 2,401 birds (Lepage, 2021) and 1,648 reptiles (Tolley *et al*., 2016) and ∼600 amphibians (Channing & Rodel, 2019)), a conservative estimate would be around 15,000 MBPs, suggesting that only ∼1% of the total African MBP diversity is currently known. Thus, pathogen and vector discovery, as well as identifying their reservoir hosts would be highly productive endeavor, especially if targeting lesser-known vertebrates and mosquitoes. The global emergence of “benign” zoonotic pathogens such as ZIKV, CHIKV, and WNV illustrate the need for comprehensive knowledge of the MBPs, including those transmitted among wild animals by unknown vectors. Targeting mosquito subgenera with a high fraction of African species that presumably had longer time to be co-opted as vectors by African pathogens, such as species of *Eretmapodites, Catageiomyia (Aedes), Neomelaniconion (Aedes), Albuginosus (Aedes), Hopkinsius (Aedes), Maillotia (Culex)*, and *Barraudius (Culex)* might yield many new MBPs. The network of mosquitoes and MBPs defined by their significant co-occurrence in the same countries (Fig. S2) identifies putative sylvatic vectors (Supp. Results and Discussion).

### The area occupied by African mosquitoes and MBPs: drivers and implications

With a quarter of the species occupying a single country, ∼50% in 1–3 countries, and only 5% or less present in >25 countries (Fig. 2), most mosquitoes and MBPs occupy relatively small ranges. Endemicity in African mosquitoes is lower than that reported globally (50%) (Foley *et al*., 2007), probably because African islands were excluded from our analysis. Based on the median total area occupied by a mosquito and a species of MBP (see Results), their typical ranges cover 10% and 7% of continental Africa, respectively. These range sizes can be approximated by squares with sides of 1,760 km and 1,500 km, respectively. The distributions of range size in African mosquitoes and MBPs are strikingly similar, although the mosquito median is larger than that of the MBPs (see Results), suggesting that most African MBPs are transmitted by one or just a few narrow-range mosquitoes in sylvatic cycles among their wild host species. MBPs with one or few mosquito vectors are probably specialized to these vector species and vertebrate hosts, whose ranges ultimately limit pathogen spread. Therefore, adapting to multiple vector species is likely a prerequisite for initiating range expansion in MBPs (see more below) and given the small number of domesticated mosquitoes (Fig. 2a). Considering that the majority of the MBPs circulate in wild vertebrates (Fig. 1c) in a relatively small area, this state might have represented the original phase of today’s most widespread MBPs (Fig. 2b), which have since undergone range expansion. This may also apply to MBPs that will emerge in the future (below).

Surprisingly, most species of *Aedes, Anopheles*, and *Culex* occupy a typically small area (1–3 countries), whereas the widespread outlier species (>30 countries) include *Ae. aegypti, Ae. vitattus, Cx. quinquefasciatus*, and ten Anopheles species, e.g., the primary malaria vectors *An. gambiae, An. arabiensis, and An. funestus*, as well as the more zoophilic *An. pharoensis, An. squamosus, An. coustani, An. ziemanni*. (Fig. 2a). These species feed preferentially on people and/or domestic animals and are well adapted to the domestic environment. Notably, at least six of these thirteen species were intercepted at high altitudes (40–290 m above ground) in the Sahel of Mali (Huestis *et al*., 2019), indicating that windborne long-range migration is a common trait of these exceptionally widespread species, as for other insects (Pedgley *et al*., 1995; Reynolds *et al*., 2006; Chapman *et al*., 2012; Drake & Reynolds, 2012). The high proportion of widespread *Anopheles* species suggests faster adaptation to domestic environment and/or increased dispersal capacity. These traits may mutually reinforce each other because the widespread presence of domestic settings minimize the risk of ending long-range migration in an inhospitable habitat.

Similarly, the exceptionally widespread MBPs whose range exceed 20 countries include only nine species (Fig. 2). Except for WNV, which is transmitted between wild birds, six are transmitted between people: PLFLC and PLVIX (>30 countries), PLOVL, DENV, YFV, and CHIKV, whereas RVFV and anthrax are transmitted between domestic animals. All these MBPs are vectored by one or more of the most widespread mosquitoes: the human plasmodia are primarily vectored by *An. gambiae, An. arabiensis, An. funestus*, (as well as by *An. coluzzii* that occupies West and parts of Central Africa) and secondarily by *An. pharoensis, An. squamosus, An. coustani*, and *An. ziemanni*. Likewise, DENV, YFV, and CHIKV are vectored primarily among people by *Ae. aegypti*, whereas RVFV is transmitted among domestic animals by several species including *Ae. vitattus, Cx. quinquefasciatus, An. pharoensis, An. squamosus*, and *An. coustani* (Linthicum *et al*., 1985; Seufi & Galal, 2010; Tantely *et al*., 2015; Braack *et al*., 2018). WNV is also transmitted by multiple mosquito vectors, including *Cx. quinquefasciatus and An. rufipes* (Braack *et al*., 2018; Ndiaye *et al*., 2018). Finally, anthrax which is only secondarily transmitted by mosquitoes (mechanically) is transmitted among domestic and wild animals primarily via ingestion and inhalation of the bacteria from soil or plants, and can be transported hundreds of kilometers by domestic or wild host animals, e.g., vultures, elephants (Purdon *et al*., 2018; Phipps *et al*., 2019). The role of mosquitoes and other flies (Turell & Knudson, 1987) in transporting anthrax should not be disregarded. Indeed, an outbreak of the mechanically transmitted myxomatosis (caused by Myxoma virus) was attributed to windborne mosquitoes flying from Australia to Woody Island, a distance of 320 km (Garrett-Jones, 1950). This observation adds to others in support of pathogens transmitted by windborne mosquitoes over long distances (Garrett-Jones, 1962; Sellers, 1980; Kay & Farrow, 2000; Reynolds *et al*., 2006; Huestis *et al*., 2019; Sanogo *et al*., 2021). These results suggest that the key features of the few exceptionally widespread MBPs include transmission among people or domestic animals, as well as adaptation to at least one of the exceptionally widespread mosquito vectors, and often to other mosquito vectors that may be important to maintain the virus in sylvatic cycles.

These results assume that the distributions of the mosquitoes and (the known) MBPs is sound. More comprehensive surveillance is expected to add distribution records and shift some of our species to the right side of the L-shaped distributions (Fig. 2). The fraction of country endemics will probably decrease, but the L-shaped distribution may remain, because many newly described MBPs of wild vertebrates with modest ranges will be added. Considering that only part of a country is actually included in most species’ ranges, accurate location data may identify these parts, thus the net change in the typical range may be modest.

#### Diversity, endemism and composition of mosquitoes and MBPs across the continent

Consistent with ample evidence on the decrease of species richness with latitude, including studies of worldwide mosquitoes and general parasites and pathogens (Guernier *et al*., 2004; Foley *et al*., 2007; Wilkerson *et al*., 2021), African mosquito and MBP diversities measured by species richness are similarly concentrated along the equatorial forest peaking in Central Africa. A secondary high “ridge” of high diversity stretches along the eastern coast from Kenya to South Africa, whereas the lowest diversities are found across North Africa (Fig. 3). Mosquito and MBP exhibit corridors of moderate species-richness along the Sahel (Mauritania to Chad), and between Central Africa and both East and Southern Africa (Fig. 3). These corridors’ continuity and association to areas of seasonal aridity - inhospitable to mosquitoes, attest that they represent natural features (below). Unlike species richness, mosquito endemicity reveals two or three hotspots, whereas surrounding countries possessed few or no endemic species (Fig. 4). Endemic mosquito species are concentrated in the Cameroon and South Africa, followed by Uganda, Kenya, Tanzania, Angola and the DRC. The African equatorial forest, which is known for its high biodiversity combines stable conditions with diverse habitats, large area, and mountains (>1,000 m above sea level) that promote speciation and accumulation of species adapted to cooler habitats that found refuge in higher elevation (Lomolino 2020). Thus, higher rates of speciation, lower rates of extinction, and high ecosystem diversity can explain the high richness and endemicity of mosquitoes, MBPs (Figs. 3 and 4), and vertebrate species (Burgess et al. 2004). Somewhat different constellations of these factors extend the East and Southern Africa around the Rift System, which can also explain its high biodiversity (Burgess et al. 2004).

Unlike the higher richness and higher endemicity region of Central Africa, the markedly low ratio of endemicity to richness across Sahelian countries (Figs. 3 and 4) suggest a fauna with high propensity for long-range migration, which allow these mosquitoes to benefit from the ephemeral habitats that provide ideal conditions during the short Sahelian wet season (Reynolds & Riley, 1988; Pedgley *et al*., 1995; Drake & Reynolds, 2012; Huestis *et al*., 2019; Florio *et al*., 2020). Indeed >40 species of mosquitoes were intercepted at high altitudes (40– 290 m above ground) in the Sahel of Mali alone, representing ∼50% of the documented mosquito fauna (Huestis et al. 2019, and Yaro et al. unpublished), as predicted around seasonal ecosystems (Southwood, 1962; Drake & Gatehouse, 1995; Florio *et al*., 2020). Conversely, the high endemicity/richness ratio in equatorial regions (Figs. 3 and 4) suggests a lower propensity for long range migration.

The landscape of endemicity of MBPs shows a focal hotspot in Central African Republic, towering over all countries (Fig. 4). This exceptional endemicity is difficult to reconcile solely by the effect of species richness and country’s area (a determinant of the latter, Fig. 4) or the high biodiversity of the equatorial forest. In part, it might be explained by biased sampling; for example, research centers on yellow fever were established over 90 years ago in Nigeria and Uganda, leading to the discovery of new viruses e.g., WNV, ZIKV, and Semliki Forest virus (SFV). Within two decades, additional virus research centers were established in South Africa, Egypt, Ivory Coast, Senegal, Central African Republic, Kenya, Tanzania, DRC, and Sudan (Rosenberg *et al*., 2013; Vasilakis *et al*., 2019). More arbovirus surveillance was carried out around these centers, leading to differences in virus diversity among countries. Regional differences in diversity cannot be reconciled with these centers because the centers were distributed across all regions. Between-region variation in surveillance effort is much smaller than that between countries, therefore, regional analysis is used to test the main country-based results. For example, the higher fraction of region vs. country endemicity (40% vs. 25%) of the mosquito and MBP species, finding 60% of the mosquitoes and MBPs in one or two regions, and only 3% (mosquitoes) and 11% (MBPs) across the continent (Fig. 5) are consistent with country-based results. Additionally, despite sharing ecozones and biomes across our regions (Burgess *et al*., 2004), the regional analysis revealed compositional heterogeneity in mosquitoes and MBPs across the continent (Figs. 5, 6). The country dendrograms based on composition of mosquitoes and MBPs generally clustered together countries of the same subdivisions (Fig. 6). Remarkably, the clustering of regions into subdivisions based on the composition of the mosquito and MBP faunas were nearly identical (Fig. 6). Similar to plant and vertebrate biogeographical results (Burgess *et al*., 2004) our sub-Saharan Africa and North Africa divisions (Fig. 6) match the Palearctic and the Afrotropical faunal realms and highlight the Sahara as a geographic barrier. Clustering West and Central Africa regions together, separately from the cluster of East and Southern Africa (Fig. 6) fits also with the vertebrate biogeographical landscapes including those of mammals and birds (Burgess *et al*., 2004). Our West and Central regions share the Sudanian, Sahelian and Equatorial (Guinean-Congolian) zoogeographical zones whereas our East and Southern Africa regions overlap with the Zambezian and South African zoogeographical zones (Linder *et al*., 2012). The high mountains along the Rift System probably contribute to the separation between the East and Central regions. The clustering of countries based on mosquito composition indicated a subdivision of our West and Central African regions into Sudano-Sahelian and Equatorial subregion as indicated by grouping of Chad with Niger and Ivory Coast with Central African Republic (Fig. 6) showing correspondence between our results and the zoogeographical zones identified by Linder *et al*. (2012). Altogether these corresponding patterns add support for a strong bio-geographical signal in our results.

As for the distribution of range area (above), the regional composition of the MBPs was nearly identical to that of the mosquitoes, raising the question “Has MBPs geographical organization been shaped primarily by their mosquito vectors or by their vertebrate hosts?” Although both hosts and vectors limit the MBPs range, there is some evidence supporting a greater role for the mosquito vectors. Both mammals and birds exhibit large areas of high species richness and endemicity in East Africa and in smaller areas in Central, West, and Southern Africa (Burgess *et al*., 2004), unlike African mosquitoes and MBPs. Likewise, the Sudano-Sahelian area from Senegal to Sudan exhibits the second highest level of African mammals endemicity (Burgess *et al*., 2004). In birds, aside from a hyper-endemic area in East Africa, the remainder of the continent is nearly devoid of endemics (Burgess *et al*., 2004) as expected from the most mobile terrestrial vertebrate class. Although salient biodiversity features of MBPs are more similar to mosquitoes than to mammals and birds, resolving this question requires data on species richness and endemicity using the same unit area, which is beyond our analysis and data.

### A model of MBD range expansion and predicting emerging MBDs of the future

The high similarity in geographic organization of African MBPs and mosquitoes (Figs. 2–6 and S1a), and the facts that most MBPs circulate in wild vertebrate hosts (Fig. 1d) within a narrow distributional range – remarkably similar to that of mosquitoes (Fig. 2) – suggest that typically, MBPs are vectored by one or few mosquito species. Unlike this ‘original state’ of African MBPs, human malarias, YFV, WNV, RVFV, ZIKAV and CHIKV represent a small subset of exceptionally widespread African MBDs, many of which have expanded across the continent and beyond. Following Wolfe and colleagues (Wolfe *et al*., 2007), the biogeographical differences observed between the ‘original’ and the ‘emergent states’ points to a plausible process leading to range expansion of MBDs. The last phase of this range expansion includes continuous transmission (i) between people or domestic animals, (ii) by vectors that feed preferentially on these hosts and are themselves exceptionally widespread, e.g., human malaria (*An. gambiae, An. arabiensis, An. funestus* among others) and chikungunya (*Ae. aegypti*). The preceding phase would include a capacity to circulate in humans or domestic animals for a single season or a few years, depending whether vector populations are perennial or seasonal (Sellers, 1980; Linthicum *et al*., 1999; Diallo *et al*., 2005; Hanafi *et al*., 2010). How rapidly does the ‘original state’ develop into the widespread ‘emergent phases’? Unlike pathogens that are directly transmitted among hosts, the dependence of MBPs on both wild vertebrate and mosquito vector species (that blood-feed preferentially on reservoir species), range expansion requires sequential adaptations to achieve transmissibility by new vectors that expand the breadth of the MBP host species. Host and vector switching typically face fitness tradeoffs linked to specialization on particular host and vector species (Hellgren *et al*., 2007; Joy *et al*., 2008; Vasilakis *et al*., 2009; Molina-Cruz *et al*., 2013b, 2013a, 2015, 2020; Ricklefs *et al*., 2017). Thus, we expect that a typical range expansion of MBP requires a longer intermediate phase than a directly transmitted pathogen. Evidence in support of such a slow process is found in the rare occasions in which avian plasmodia (and other haemosporidia) found in migrant birds could be established in resident birds in both the Northern and Southern Hemispheres (Hellgren *et al*., 2007; Ricklefs *et al*., 2017). During the intermediate phase, the number of vectors and host species slowly increase, facilitating a gradual increase in the geographical range of the MBP. Once the MBP attains transmissibility into and from human or domestic animals by at least one of the domesticated vectors, the transition into the last phase is complete and a rapid final range expansion is expected worldwide.

Accordingly, a larger than typical geographical range is a marker of a MBP in the intermediate phase (above), in which the MBP has expanded its vector and/or vertebrate range. Except for small coastal ecozones, the African biogeographical units are typically wider across their east-west axis than across their north-south axis (Burgess 2004), suggesting that the longer the north-south dimension of a MBP’s range, the more likely it is to be transmitted by multiple vector species across multiple host species. Hence, we propose that MBP total range size, estimated as the sum of the area (of the countries) in their range and maximal north-south length of their range be used to gauge its range expansion phase. Whereas these range size measures are expected to be largest in MBPs circulating among humans and domestic animals and smallest for those circulating in wild mammals, it is less clear if MBPs circulating in wild birds are larger than those of wild mammals. Both measures were larger for MBPs circulating in humans and domestic animals for the median and the 75^th^ quantile (area: P<0.0001, t>8 df=116, north-south: P<0.0001, t>5 df=116, Figs. 7a, 7b), but no significant differences were found between MBPs circulating in mammals and birds even using one side test (area: P>0.31, t<0.11 df=116, north-south: P>0.11, t<1.2 df=116, Figs. 7a, 7b). Contrary to our expectation, the north-south distance seems to “saturate” faster than the total area (Fig. 7), indicating that it may be more sensitive to early range expansion than to later stages of MBD range expansion. The total area of most MBPs transmitted among humans or domestic animals (undergone range expansion, N=15) cover area 10–20 ×10^6^ km^2^ and their north-south distance spans 40-60°, whereas the majority of MBPs (>90) cover <4 ×10^6^ km^2^ and north-south distance >15° (Fig. 7c). Except eight MBPs transmitted among wild birds that have a long north-south distance, 30 MBPs occupy intermediate ranges covering area 4–10 ×10^6^ km^2^ and north-south distance of 20–50° (Fig. 7c). Accordingly, this group is enriched with species that are currently at the intermediate phase of range expansion. This model enables identification of putative MBPs with elevated risk for range expansion such as Usutu (USUV), Wesselsbron (WSLV), Akabane (AKAV), Spondweni virus (SPOV), and ONNV, which are elevated for both measures (Fig. 7c). This approach putatively identifies expanding pathogens during the intermediate phase of range expansion even before they infect humans or domestic animals. Monitoring changes in geographical range as well as the MBP host range and vector range would be key to evaluating these aspects of disease emergence. Validating this biogeographical ranking with independent risk predictions will increase confidence in the subset of MBPs of elevated risk. For example, the number of vectors and hosts in which a pathogen is found in and the numbers it can be transmitted from may be used as independent markers of the MBP’s prospects to undergo further range expansion. Experimental evidence about the pathogen compatibility and capacity for transmission e.g., (Haddow *et al*., 2016) with the most widespread vectors and domestic hosts (Fig. 2), will further augment the risk assessment. Development and testing of such models will advance understanding and predictive capacity of range expansion as a component of disease emergence. Evaluation of the pathogenicity and impact that a MBP would have on human and domestic animals are beyond the scope of this analysis, but the possibility of increased virulence linked to transmissibility in these new hosts by domesticated vectors, e.g., ZIKV – should not be ignored.

**Figure 7.**
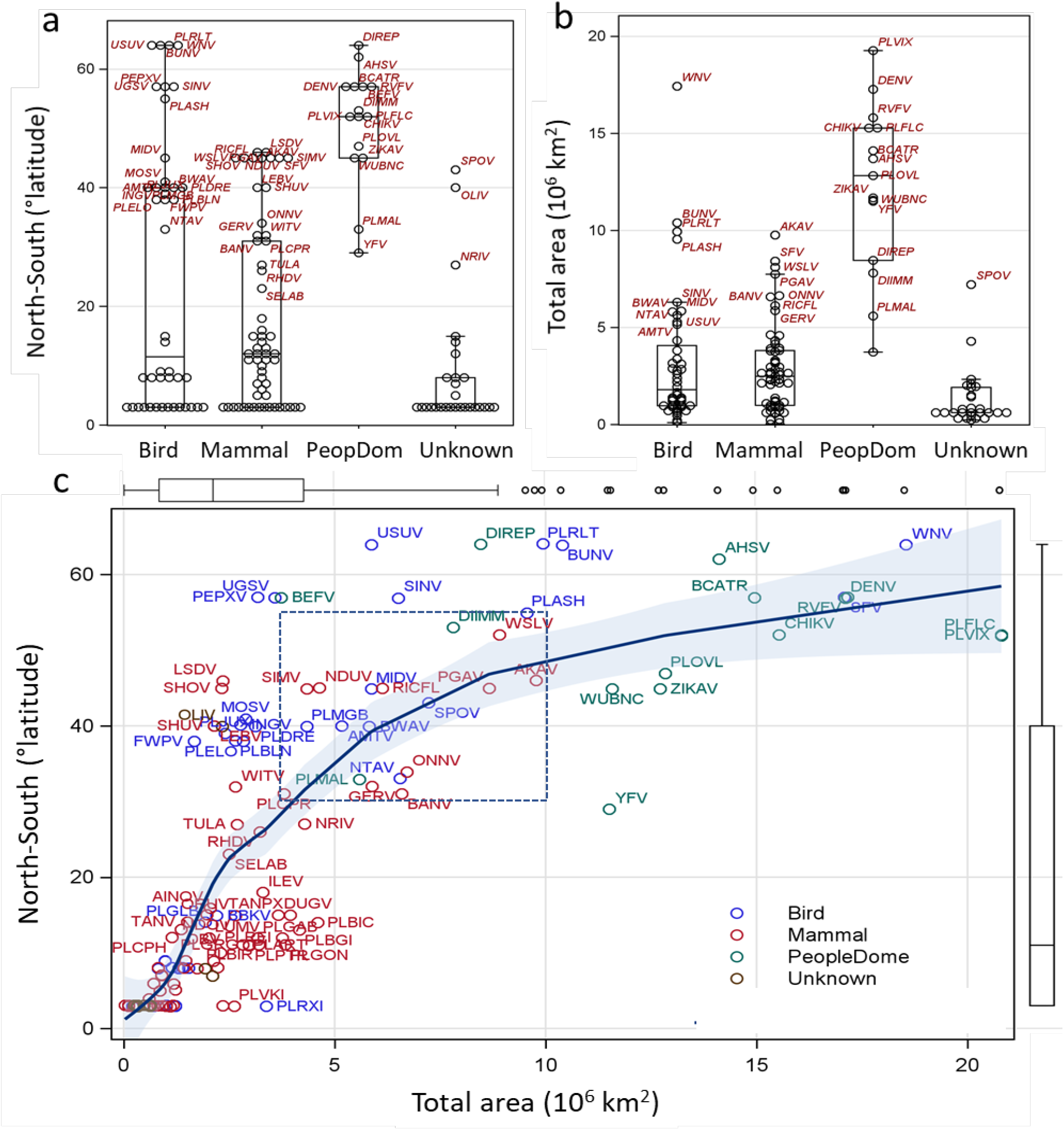
Ranking of African mosquito-borne pathogens by their range area and maximum north-south distance to estimate their phase of range expansion. a) Variation between host groups in north-south distance (latitude degrees) and b) Variation between host groups total range area (10^6^ km^2^). c) relationship between MBP’s total area (10^6^ km^2^; X-axis) and the maximum north-south distance (degrees latitude; Y-axis) using local regression (loess and 95% CLM) on all MBDs (N= 150). Box plots along axes display distributions of corresponding variables. Acronym of MBPs are given for those with total area larger than 2.5 or north-south longer than 5° and colors denote host group (birds were used if birds and mammals are thought to act as natural hosts). Box draws attention to putative MBPs at intermediate phase of range expansion (excluding MBPs of birds and domestic animals, see text).

## METHODS

The database and our analysis refer to continental Africa (surrounded by the Mediterranean Sea to the north, the Indian Ocean to the east and the Atlantic Ocean to the west), excluding all islands (e.g., Cape Verde, Comoros, Madagascar, Mauritius, Seychelles, São Tomé and Príncipe) because island biogeography requires consideration of multiple factors, such as distances to the nearest mainland and to other islands, historical formation of the island, existence of past terrestrial bridges, etc., which deserve a separate treatment. Very few records of mosquitoes and MBPs can be found for Eswatini, Lesotho, South Sudan and Western Sahara. Moreover, parts of their records are included in their previous political affiliations, e.g., South Sudan in Sudan. Therefore, these countries are not listed in our analysis; instead, our analysis, pertains to 45 countries, with few countries that subsumed those in the past and still “contain” their records, e.g., “Sudan and South Sudan” being used (Table S1). Because countries differ in surveillance effort, grouping neighboring countries into regions minimizes variation in surveillance effort variability and was used to test country-based patterns. Unlike the geopolitical regions with the same names, our five regions were defined to maximize distances among regions, accommodate latitudinal variation, and minimize inter-region enclaves (Fig. 5).

Our African mosquito distribution data (Supp. File 1) was initially generated based on the global distribution lists, updated to 2017 (Wilkerson *et al*., 2021). We updated records of anophelines in sub-Saharan countries (Irish *et al*., 2020), and culicines following country-specific lists recently published for Mali (Tandina *et al*., 2018), Mauritania (Lemine *et al*., 2017), Morocco (Trari *et al*., 2017), and incorporated records for southern African countries (Jupp, 1996). Information on global diversity of mosquitoes was recently updated (Wilkerson *et al*., 2021) and allowing reconciliation of species identifications that were later revised, e.g., *Culex tigripes* and *Lutzia tigripes* or *An. arabiensis* and *An. gambiae*. Subspecies were not included in our data.

The mosquito-borne pathogen (MBP) distribution data was generated based on hundreds of references listed in Supp File 2, providing they met the three criteria as follows: A peer-reviewed scientific source (or a source, e.g., the CDC arbovirus catalogue, listed in peer-reviewed sources) reported that the MBP has been i) naturally transmitted in continental Africa, ii) to a terrestrial vertebrate host, iii) by mosquito vector, to the extent that this mode of transmission is recognized to have an epidemiological role, even if other mode(s) of transmission play a greater role. Our database includes information whether mosquito role in the MBP transmission is secondary or primary and whether it is biological or mechanical. Strains or any sub-species definitions were not included in our analysis. To ascertain accuracy of our MBP records, we compared our data with the CRORA database (*Centre de Référence OMS sur la Recherche des Arbovirus et des Fièvres Hémorragiques (CRORA): www.pasteur.fr/recherche/banques/CRORA* (discontinued since 2015) and the EID2 database (Wardeh *et al*., 2015) (as of September 2021) among other sources. Only records that met our above criteria were included in our database. By confining our records to continental Africa, the term endemic refers to a species found in one African country (or region, when specified), however, although uncommon, the species may be also found outside continental Africa.

Information on land mass of the World and of continental African countries (The-World-Bank, 2021) were used to calculate the proportion of area of continental Africa from the land worldwide and total area per species. Accordingly, the total area of the worldwide and continental Africa we used are 148,568,946.1 and 296,63,582.0. Global coordinates central position of each African country (Google Developers, 2021) were used to computed maximum north-south range distances for each MBP.

### Data analysis

Goodness of fit χ^2^ tests implemented by Proc Freq (SAS Institute, 2012) were used to assess if diversity in a particular area was higher than predicted by the relative size of the area. Exact tests were used if expected values were smaller than 5. Confidence intervals (distribution free) of medians were computed using Proc Univariate (SAS Institute, 2012) based on order statistics (ranks). Person correlation, linear, and quadratic regression models to relate biodiversity measures with country area were implemented by Proc Reg (SAS Institute, 2012). Quantile regression implemented by Proc Quantreg (SAS Institute, 2012) extends the general linear model for estimating conditional change in the response variable across its distribution as expressed by quantiles, rather than its mean (though the median is similar to the mean in symmetric distributions). It does not assume parametric distribution (e.g., normal) of the random error part of the model, thus it is considered semiparametric. The value of this analysis is that it allows us to address variation among the medians of various groups and also across quantiles even when the mean and the median are unchanging. The parameters estimates in linear quantile regression models are interpreted as in typical general linear models, as rates of change adjusted for the effects of the other variables in the model for a specified quantile (Cade and Noon 2003). We used matrices of presence absence of mosquitoes or MBPs to compute matrices of Jaccard distances between regions or countries (separately), using Proc Distance (SAS Institute, 2012) and used the Ward method in Proc Cluster with height measured by R^2^ (the proportion of variance accounted by the clusters) to produce and plot dendrograms.

## Acknowledgements

We thank Drs. Roy Faiman, Alvaro Molina-Cruz, Jose’ MC Ribeiro (NIH/NIAID), Don R. Reynolds (University of Greenwich, UK), Tom Burkot (James Cook University, Australia), Phil Lounibos (University of Florida) for providing helpful comments based on earlier versions of this manuscript and for exciting discussions. This study was supported by the Division of Intramural Research, National Institute of Allergy and Infectious Diseases, National Institutes of Health, Bethesda MD. Y-ML was supported by the Global Emerging Infections Surveillance Branch of the Armed Forces Health Surveillance Division (AFHSD-GEIS) FY21 project P0030_21_WR. The views, opinions and/or findings contained in this report are those of the author(s) and should not be construed as an official U.S. Department of the Army, Department of the Army position, policy or decision unless so designated by other documentation.

## Supplemental Information

Supplemental Results and Discussion

Table S1: Genera and subgenera of mosquitoes in continental Africa

Table S2: African countries included in this analysis: area and centroid coordinates

Supp. Data File 1: African mosquito faunal list per country (to be included with publication in refereed journal)

Supp. Data File 2: African mosquito-borne pathogens: Acronym, taxonomical affiliation, transmission mode, and distribution by country, with references (to be included with publication in refereed journal)

## Supplemental Results and Discussion

The similarity in species richness of MBPs and mosquitoes (Fig. 3) is also expressed by the high positive correlation coefficient between these indices (r= 0.76, N= 45, P< 0.001 Fig. S1a), although the correlation in species endemicity was far lower (r= 0.26, N= 45, P= 0.09, Fig. S1b). Because conventional pathogen detection requires species-specific diagnostic test that have been developed for common and widespread pathogens, endemic pathogens are expected to be under-detected. Furthermore, this weak correlation may also reflect sampling effort inequality of uncommon MBPs among countries (see Main text).

Although the overall regional distributions of mosquitoes and MBPs are very similar (Fig. 5), the differences (Fig. S1c) reveal a higher fraction of cross-Sahara MBPs (found in all five regions), whereas a higher fraction of mosquitoes if found across sub-Saharan Africa (P< 0.01, Fig. S1c). This pattern suggests that relatively few MBPs, albeit more than mosquitoes, are transported by their vertebrate host(s) across the Sahara. Additionally, MBPs that have been recently introduced into Africa, eg., DENV, may have been arrived into multiple parts of the continent, and being already adapted to the domestic environment, may have spread rapidly. Unlike pathogens that are easily transported by human and domestic animals, fewer mosquitoes represent this subgroup, e.g., *Ae. albopictus* and *An. stephensi*.

**Fig. S1.**
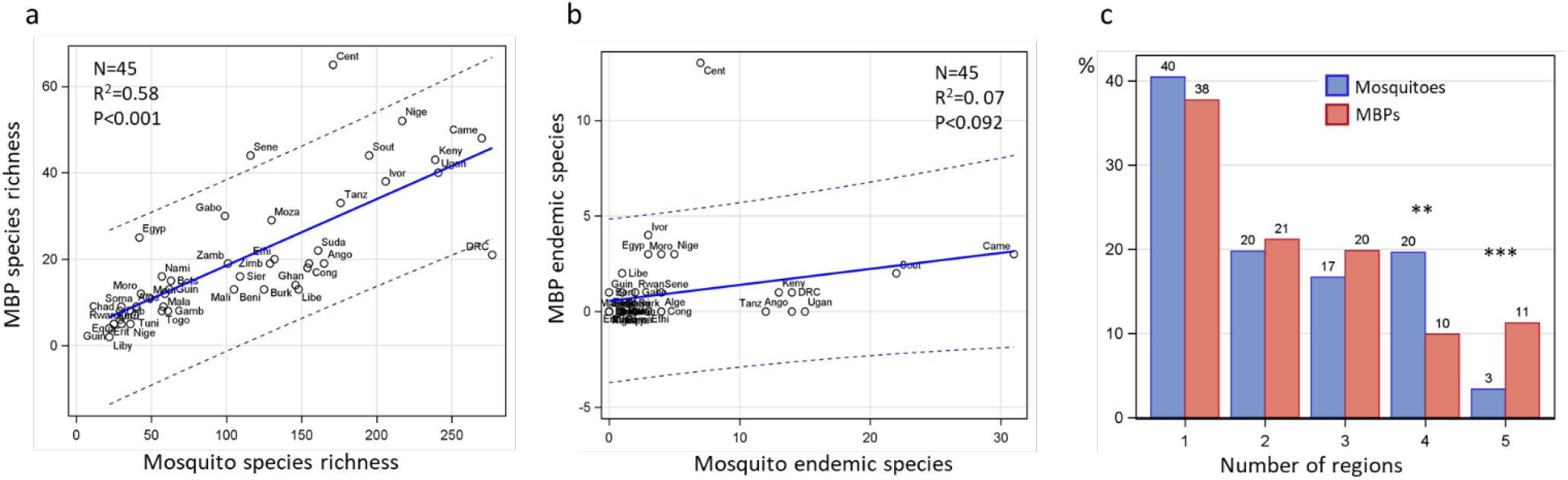
Congruence between MBPs and mosquitoes in country diversity (a), endemicity (b) and in the number of regions occupied (c). The relationship between MBPs and mosquitoes in species richness (a) and endemicity (b) is shown using a linear regression (blue solid line) and 95% CI for individual observations (countries, dotted line) with corresponding statistics. Names of countries are abbreviated to the first four letters. c) Differences in the distribution of the number of regions occupied by mosquitoes and MBPs in percent (above bars). Statistically significant differences (determined using 2×2 heterogeneity χ^2^ tests) are shown by asterisks: ** and *** denote P<0.01 and P<0.001, respectively.

Mechanical transmission of MBPs by mosquitoes is usually disregarded by vector biologists, because it is not the primary mode of pathogen transmission (Fig. 1c), which perpetuates this attitude despite limited information about it. In Africa, only pox viruses and bacteria are reported to be transmitted mechanically by mosquitoes (Fig. 1c). The epidemiological contribution of mosquito transmission of these MBPs in short- and long-range spread of the pathogens is poorly known (but see main text), as well the extent of the vector range used by these pathogens. Further study and surveillance of pathogens transmitted mechanically by mosquitoes (especially bacteria) would reveal new grounds.

### Geographical assemblages of mosquitoes and MBPs

Species that co-occur more than expected by chance define regional assemblages that can underlie similar ecological preferences or co-dependence. We used Veech co-occurrence index (Veech, 2013) to evaluate which pairs of species co-occur positively across countries (joint country occurrence is higher than expected by chance), negatively (joint country occurrence is lower than expected by chance), or randomly (joint country occurrence is not different than expected by chance). Considering the whole continent (45 countries), the results revealed that 73% of the mosquito pairs were random, 24% were positive (P<0.05), and 3% were negative (P<0.05, N= 57,847 testable species pairs). North Africa might have inflated significant associations because only 5% of species are found across the Sahara. Excluding the North African countries, reduced the fraction of significant associations: 19% (10,826) positive and 1% (650), negative (80% showing random associations, N= 57,326 species pairs, Fig. S2a: inset) across 40 countries. Negative co-occurrence in mosquitoes were especially common in *Anopheles* (78%) and *Culiseta* (6%), both within and across genera, whereas positive co-occurrences between species pairs were distributed across all genera (not shown). Such negative association suggests adaptive speciation in distinct environments, where assemblages are unique, and therefore not overlapping in species composition. *Anopheles* and *Culiseta* have the highest fraction of African species among the genera (except *Eretmapodites*, which is mostly equatorial; see Results), suggesting extensive speciation in Africa and distinct environment that supports this explanation.

Considering MBPs in the whole continent, 17.4% (315) of the pairs were positive and 0.2% (4) were negative out of a total of 1,807 testable pairs (Fig. S2b: inset). Because 17% of species are found across the Sahara, rather than the corresponding 5% of the mosquitoes (above), North African countries were included in subsequent analyses. Unlike the negative co-occurring MBP pairs, the number of positive co-occurring MBP pairs is far higher than that expected by chance (2.5%).

To visualize the organization of mosquito assemblages, defined by the co-occurrence analysis, positively co-occurring pairs of narrow-range species (5-10 countries) were included in a network consisting 119 pairs (Fig. S2a). The mosquito network exhibited four disjointed components, reflecting distinct assemblages: (i) Western-Central Africa cluster represented by *Cx. subrima* (main) with an equatorial cluster represented by *Ae. abnormalis*, (ii) Southern-East Africa represented by *An. confusus*, (iii) Southern-Central Africa, represented by *An. walravensi*, and (iv) a small widespread assemblage, represented by *Ae. capensis* (Fig. S2a).

The network of 53 MBPs pairs consisting of 25 species, (5-10 countries) presented a single component (Fig. S1b). While more densely connected than the mosquito network (density: 0.177 vs. 0.059), distinct yet partly-overlapping assemblages are found in Southern-East and Central Africa represented by Bwamaba/Ndumu viruses (Fig. S2b, BWAV/NDUV), showing resemblance to the Southern-East Africa represented by *An. confusus*, a smaller equatorial assemblage represented by (ORUV), resembles the mosquito cluster represented by *Ae. abnormalis*, whereas the sub-Saharan MBP assemblage represented by Middleburg virus (MIDV) resembles the small, yet widespread mosquito assemblage represented by *Ae. capensis*.

**Figure S2.**
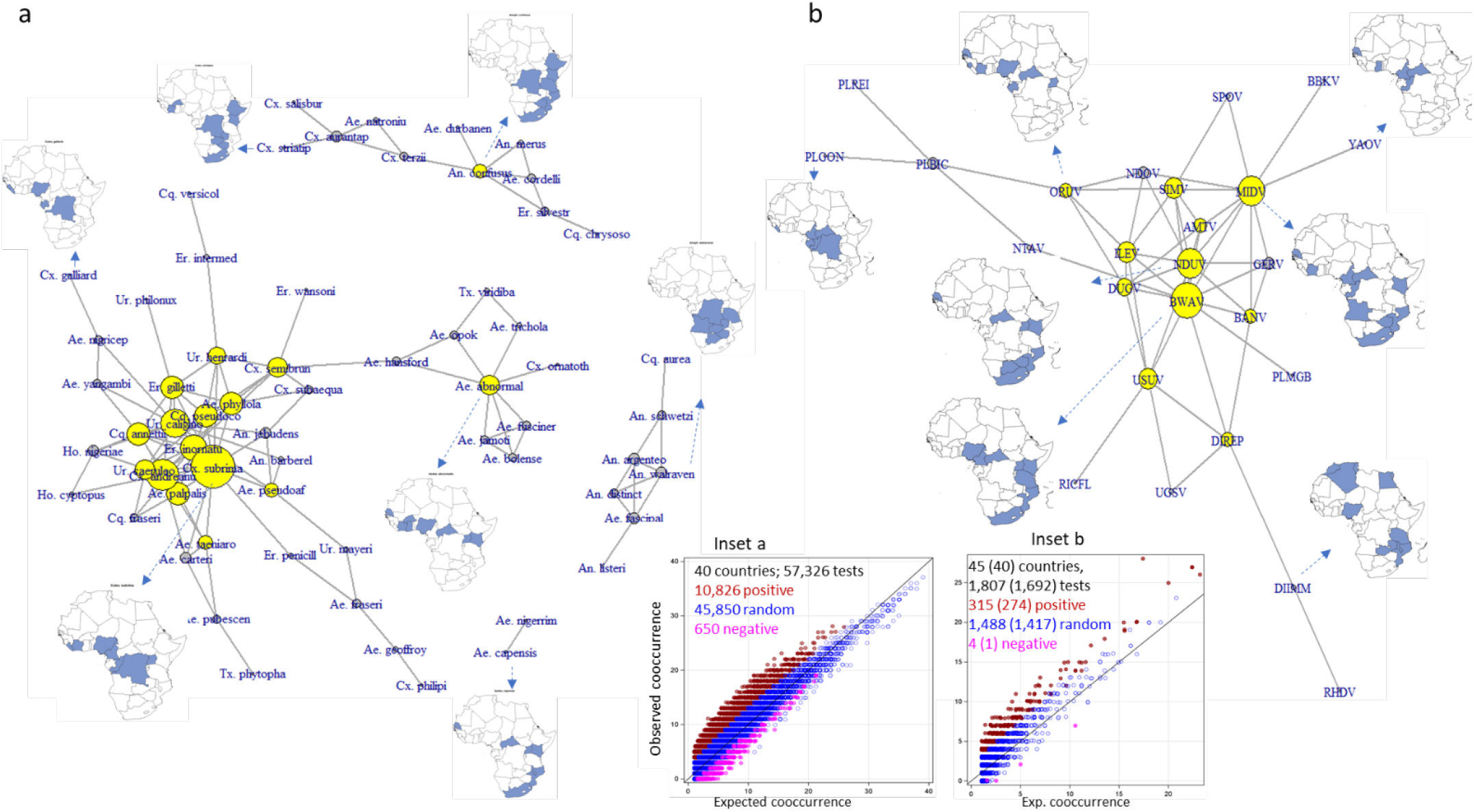
Network of positively co-occurring pairs of mosquito (a) and MBP (b) species (P<0.05), whose range is up to 10 countries (text). a) Node size proportional to its degree centrality (number of species it is significantly co-occurring with) yellow denotes >4 degrees). Distribution maps are drawn for species (broken arrows) with highest degree centrality within a component (cluster) as well as for arbitrary low-centrality species. Although North Africa was not included in the calculations of the co-occurrence for mosquitoes (text), it is included in the maps. Insets show the relationship between observed and expected joint co-occurrence (number of countries both species co-occur at) for mosquitoes (inset a) and MBPs (inset b). Positive pairs co-occur in more countries (observed) than expected denoted by the diagonal line and negative pairs co-occur in less countries than expected. Significantly positive and negative pairs are shown in red and magenta, respectively; nonsignificant pairs (random) are shown in blue.

Because mosquito transmission is the primary route of vertebrate infection in >90% of MBPs (Fig. 1c), high vector-specificity is expected for sylvatic vectors, except for the poxviruses and the three bacteria that rely on mechanical transmission. A bipartite network of mosquitoes and MBPs based on the Veech index can reveal assemblages of mosquitoes and pathogens, which might be different than their within-group assemblages. Albeit based on geography alone, linking mosquitoes and MBPs may also help identify subset of putative sylvatic mosquito vectors, which can be further refined applying other selective criteria. Veech co-occurrence analysis (over the whole continent) revealed that 83% of the testable pairs were random, 16% were positive and 1% were negative at P<0.05 (N= 21,342 testable mosquito-MBP pairs). Considering only highly positively co-occurring pairs (P<0.01; note: 2.5% of the total tests at each side are expected by chance), a bipartite network comprising of 30 MBPs and 80 mosquitoes with 194 links was plotted (Fig. S3). To simplify interpretation, mechanically-transmitted MBPs and non-blood-feeding mosquitoes were excluded and only pairs in which the pathogen joint co-occurrence was fully subsumed in that of the mosquito were retained. The largest MBP-mosquito assemblages were from Central Africa: *Plasmodium gonderi* (PLGO with 19 mosquitoes), *Pl. reichenowi* (PLRE), followed by that in Southern-East and Central Africa Bwamaba virus (BWAV, Figs. S2, S3).

**Figure S3.**
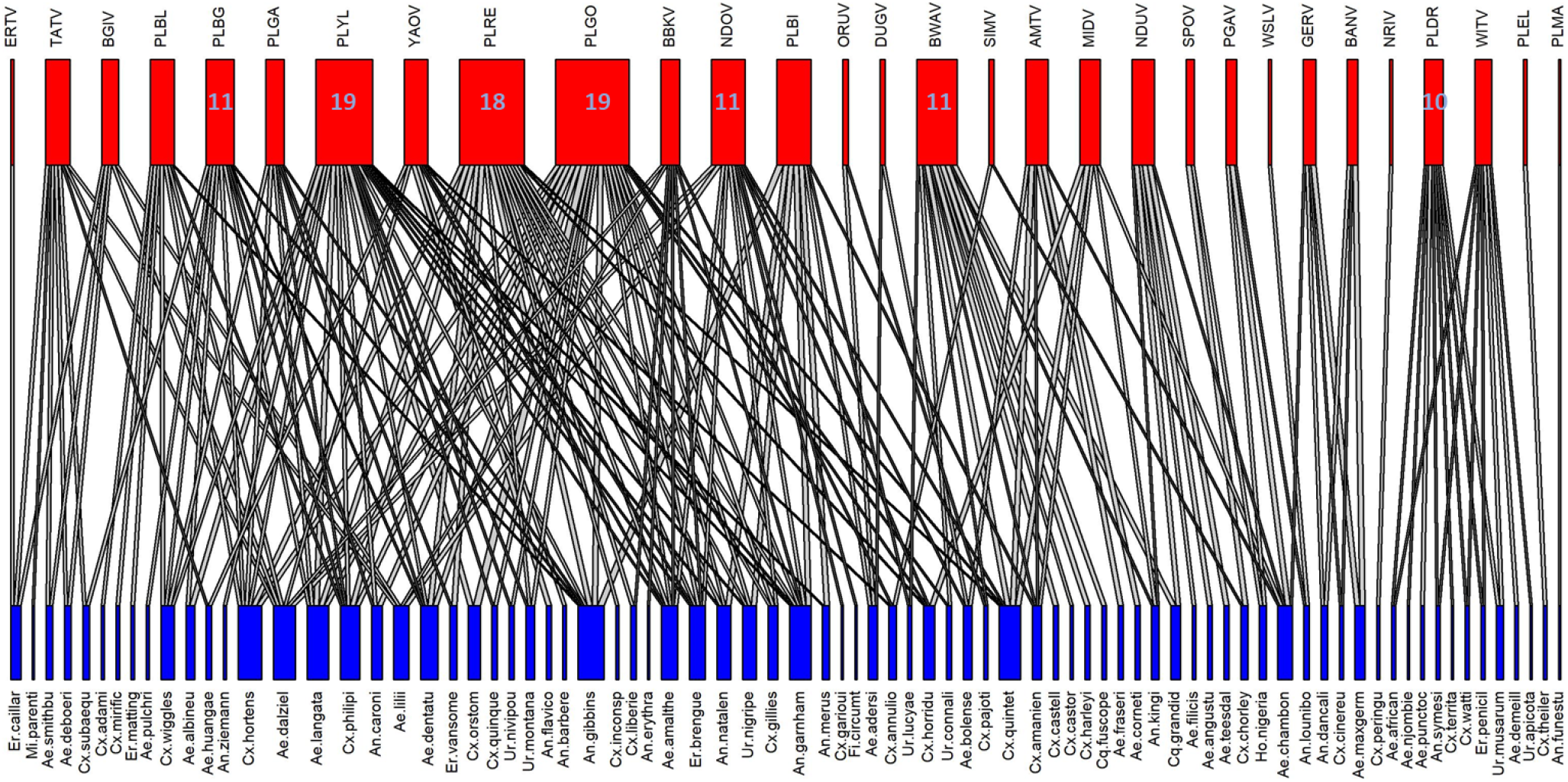
Bipartite network of positively co-occurring pairs (P<0.01) of MBP (N= 30, top-red) and mosquito (N= 80, bottom-blue) to identify putative sylvatic vectors (see text). Note: mechanically-transmitted MBPs and non-blood-feeding mosquitoes were excluded.

This network includes broad-ranging MBPs, e.g., *Pl. malariae*, whose range is subsumed only in that of *An. funestus* (Fig. S3). On average each MBP is linked to 6.5 mosquitoes (median= 6). Because geographical overlap in MBP and mosquito distribution is the only basis for linking them, the matrix included 5 non-bloodfeeding mosquitoes (*Ml. fraseri, Ml. moucheti, Tr. aeneus, Tr. viridibasis*, and *Tr. wolfsi*, not shown). Likewise, the links between *Pl. reichenowi*, which is thought to be transmitted exclusively by *Anopheles* mosquitoes also include 15 culicines. Therefore, incorporating additional criteria such as *Anopheles* to filter among putative vectors of a mammalian *Plasmodium* species, incorporating information on bloodmeal host range, and permitting partial range overlap can produce a more accurate list of putative mosquito vector species for surveillance and vectorial competence experiments. This approach can help identifying the sylvatic vectors of many pathogens.

## Supplemental Methods

The probabilistic model of species co-occurrence (Veech, 2013) was used to classify pairs of species co-occurrence as negative, positive or random based on the probabilities that two species would co-occur at a frequency less than or greater than the observed frequency if the two species were distributed independently of one another among sites. Veech index is based on an analytic probabilistic model using combinatorics to obtain the probability that two selected species co-occur at any given number of sites among those sampled. The calculations were coded in SAS (SAS Institute, 2012). Given the number of countries (sites) used in our data (40– 45), significant pairs could be classified at the P<0.05 significance level, only if both ranges cover at least 4 countries. We used Proc Gmap (SAS Institute, 2012) to generate maps. Network plots and statistics were produced using the packages igraph (Csardi & Nepusz, 2006) and bipartite (Dormann *et al*., 2008) in R (R Core Team, 2016).

**Table S1.**
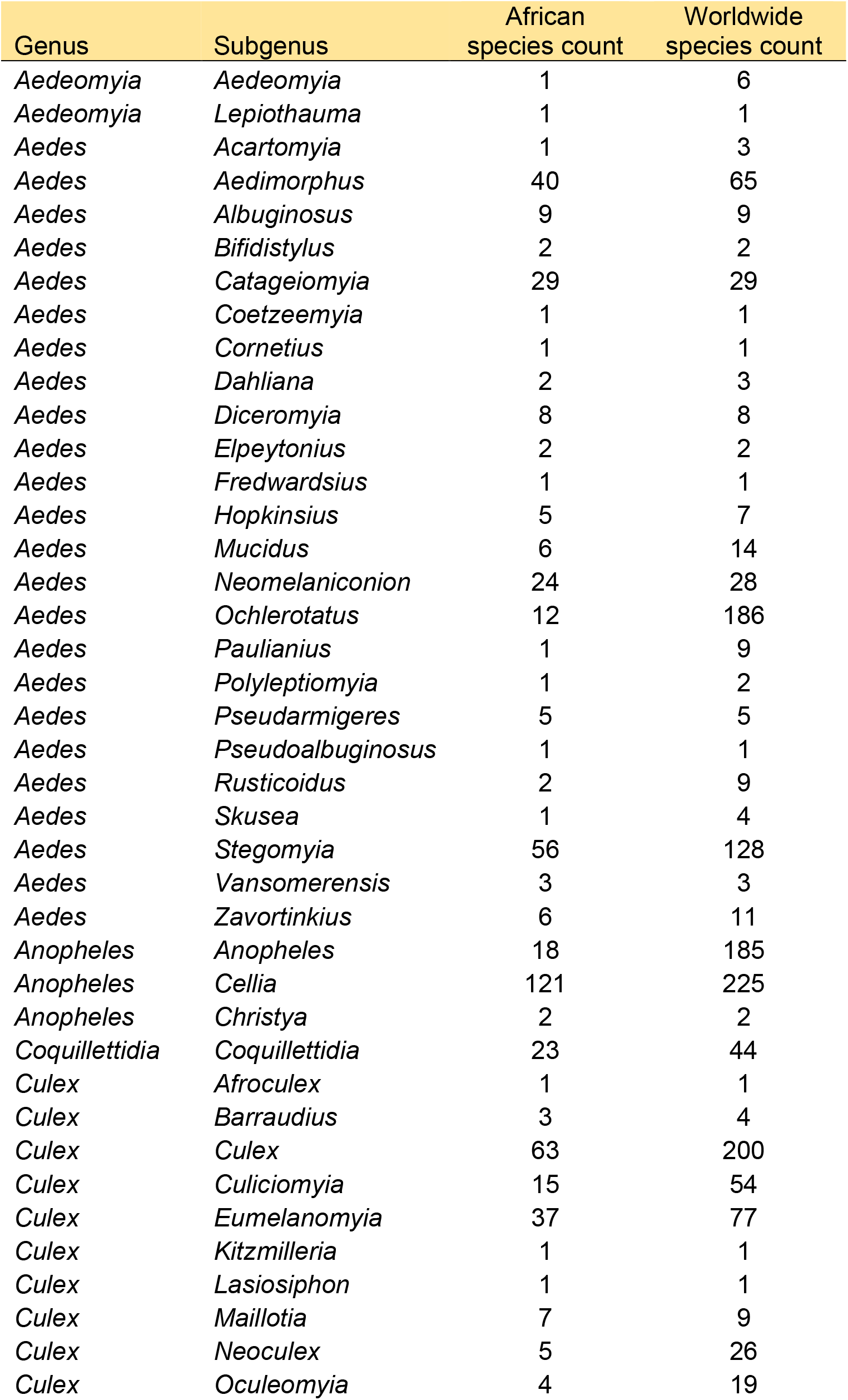
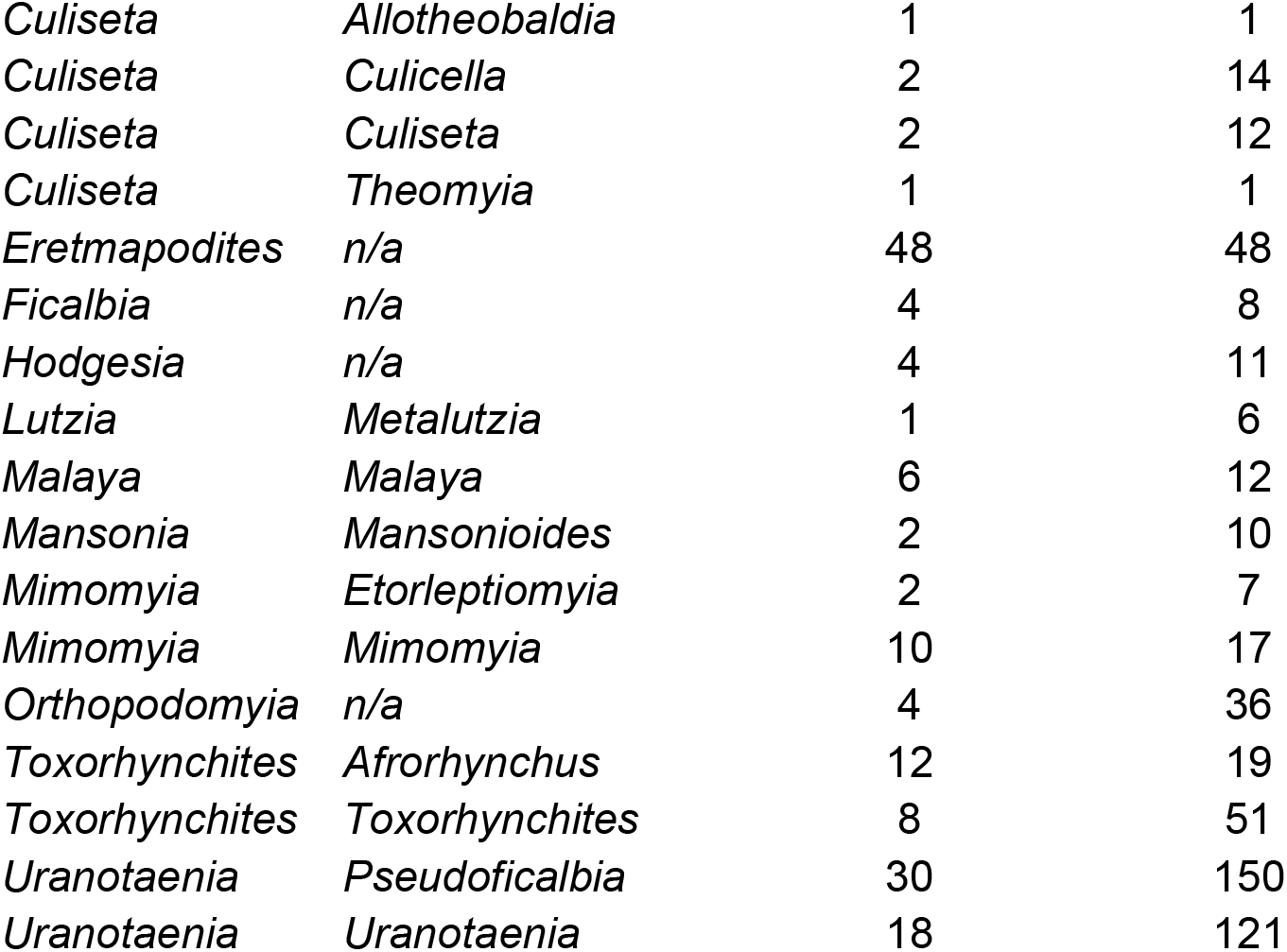
Subgenera of mosquitoes in continental Africa with their number of species in the continent and worldwide (Wilkerson *et al*., 2021)

**Table S2.** African countries included in this paper (N= 45), area, centroid position, and additional territories subsumed as used in this paper (Methods).

| Country | Area<br>(km<sup>2</sup>) | Latitude | Longitude | Subsumes |
| --- | --- | --- | --- | --- |
| Algeria | 2381741 | 28.033886 | 1.659626 |  |
| Angola | 1246700 | -11.202692 | 17.873887 |  |
| Benin | 112622 | 9.30769 | 2.315834 |  |
| Botswana | 581726 | -22.328474 | 24.684866 |  |
| Burkina Faso | 274000 | 12.238333 | -1.561593 |  |
| Burundi | 27830 | -3.373056 | 29.918886 |  |
| Cameroon | 475442 | 7.369722 | 12.354722 |  |
| Central African Republic | 622984 | 6.611111 | 20.939444 |  |
| Chad | 1284000 | 15.454166 | 18.732207 |  |
| DRC | 2344858 | -4.038333 | 21.758664 |  |
| Djibouti | 23200 | 11.825138 | 42.590275 |  |
| Egypt | 1001449 | 26.820553 | 30.802498 |  |
| Equatorial Guinea | 28051 | 1.650801 | 10.267895 |  |
| Eritrea | 117600 | 15.179384 | 39.782334 |  |
| Ethiopia | 1104300 | 9.145 | 40.489673 |  |
| Gabon | 267668 | -0.803689 | 11.609444 |  |
| Gambia | 10380 | 13.443182 | -15.310139 |  |
| Ghana | 238534 | 7.946527 | -1.023194 |  |
| Guinea | 245857 | 9.945587 | -9.696645 |  |
| Guinea-Bissau | 36125 | 11.803749 | -15.180413 |  |
| Ivory Coast | 322460 | 7.539989 | -5.54708 |  |
| Kenya | 580367 | -0.023559 | 37.906193 |  |
| Liberia | 111369 | 6.428055 | -9.429499 |  |
| Libya | 1759540 | 26.3351 | 17.228331 |  |
| Malawi | 118484 | -13.254308 | 34.301525 |  |
| Mali | 1240192 | 17.570692 | -3.996166 |  |
| Mauritania | 1030700 | 21.00789 | -10.940835 |  |
| Morocco | 710850 | 31.791702 | -7.09262 | Western Sahara |
| Mozambique | 801590 | -18.665695 | 35.529562 |  |
| Namibia | 825418 | -22.95764 | 18.49041 |  |
| Niger | 1267000 | 17.607789 | 8.081666 |  |
| Nigeria | 923768 | 9.081999 | 8.675277 |  |
| Republic of Congo | 342000 | -0.228021 | 15.827659 |  |
| Rwanda | 26798 | -1.940278 | 29.873888 |  |
| Senegal | 196723 | 14.497401 | -14.452362 |  |
| Sierra Leone | 71740 | 8.460555 | -11.779889 |  |
| Somalia | 637657 | 5.152149 | 46.199616 |  |
| South Africa | 1221037 | -30.559482 | 22.937506 | Eswatini and Lesotho |
| Sudan & South Sudan | 2505813 | 12.862807 | 30.217636 | Sudan and South Sudan |
| Tanzania | 945203 | -6.369028 | 34.888822 |  |
| Togo | 56785 | 8.619543 | 0.824782 |  |
| Tunisia | 163610 | 33.886917 | 9.537499 |  |
| Uganda | 236040 | 1.373333 | 32.290275 |  |
| Zambia | 752614 | -13.133897 | 27.849332 |  |
| Zimbabwe | 390757 | -19.015438 | 29.154857 |  |

## References

Arisue, N., Honma, H., Kume, K. & Hashimoto, T. (2021) Progress in understanding the phylogeny of the Plasmodium vivax lineage. Parasitology international, 87.

Bensch, S., Hellgren, O. & Perez-Tris, J. (2009) MalAvi: a public database of malaria parasites and related haemosporidians in avian hosts based on mitochondrial cytochrome b lineages. Molecular Ecology Resources, 9, 1353–1358.

Binder, S., Levitt, A.M., Sacks, J.J. & Hughes, J.M. (1999) Emerging infectious diseases: Public health issues for the 21st century. Science, 284, 1311–1313.

Braack, L., Gouveia De Almeida, A.P., Cornel, A.J., Swanepoel, R. & Jager, C.De. (2018) Mosquito-borne arboviruses of African origin: Review of key viruses and vectors. Parasites and Vectors, 11, 29.

Burgess, N.D., Hales, J.D., Underwood, E., Dinerstein, E., Olson, D., Itoua, I., et al. (2004) Terrestrial eco-regions of Africa and Madagascar: A conservation assessment. World Wildlife Fund, Unitied States, Island Press, Washington D.C.

Burgin, C.J., Colella, J.P., Kahn, P.L. & Upham, N.S. (2018) How many species of mammals are there? Journal of Mammalogy, 99, 1–14.

Burke, D. (1998) The evolvability of emerging viruses. In Pathology of Emerging Infections 2 (ed. by Nelson, A.M. & Horsburg, R.C.J.). American Society for Microbiology, Washington, DC, USA, pp. 1–12.

CDC. (2019) Arbovirus Catalog [WWW Document]. URL https://www.n.cdc.gov/arbocat/VirusBrowser.aspx [accessed on.

Channing, A. & Rodel, M.-O. (2019) Field Guide to the Frogs & Other Amphibians of Africa. Penguin Random House South Africa.

Chapman, J.W., Bell, J.R., Burgin, L.E., Reynolds, D.R., Pettersson, L.B., Hill, J.K., et al. (2012) Seasonal migration to high latitudes results in major reproductive benefits in an insect. Proceedings of the National Academy of Sciences, 109, 14924–14929.

Collins, W.E. & Jeffery, G.M. (2005) Plasmodium ovale: Parasite and disease. Clinical Microbiology Reviews, 18, 570–581.

Csardi, G. & Nepusz, T. (2006) The igraph software package for complex network research. InterJournal, Complex Sy, 1695.

DaMassa, A.J. (1966) The role of Culex tarsalis in the transmission of fowl pox virus. Avian Diseases, 10, 57.

Daron, J., Boissière, A., Boundenga, L., Ngoubangoye, B., Houze, S., Arnathau, C., et al. (2021) Population genomic evidence of Plasmodium vivax Southeast Asian origin. Science advances, 7.

Diagne, M.M., Ndione, M.H.D., Gaye, A., Barry, M.A., Diallo, D., Diallo, A., et al. (2021) Yellow Fever Outbreak in Eastern Senegal, 2020–2021. Viruses 2021, Vol. 13, Page 1475, 13, 1475.

Diallo, D., Sall, A.A., Buenemann, M., Chen, R., Faye, O., Diagne, C.T., et al. (2012) Landscape ecology of sylvatic chikungunya virus and mosquito vectors in southeastern senegal. PLoS Neglected Tropical Diseases, 6, 1–14.

Diallo, D., Sall, A.A., Diagne, C.T., Faye, O., Faye, O., Ba, Y., et al. (2014) Zika virus emergence in mosquitoes in Southeastern Senegal, 2011. PLoS ONE, 9.

Diallo, M., Nabeth, P., Ba, K., Sall, A.A., Ba, Y., Mondo, M., et al. (2005) Mosquito vectors of the 1998-1999 outbreak of Rift Valley Fever and other arboviruses (Bagaza, Sanar, Wesselsbron and West Nile) in Mauritania and Senegal. Medical and Veterinary Entomology, 19, 119–126.

Dormann, C., Gruber, B., interaction, J.F.-& 2008, undefined. (2008) Introducing the bipartite package: analysing ecological networks. biom.uni-freiburg.de, 8.

Drake, V.A. & Gatehouse, A.G. (1995) Insect migration: tracking resources through space and time. Cambridge University Press, New York.

Drake, V.A. & Reynolds, D.R. (2012) Radar entomology : observing insect flight and migration. CAB International., Wallingford, UK.

Epelboin, Y., Talaga, S., Epelboin, L. & Dusfour, I. (2017) Zika virus: An updated review of competent or naturally infected mosquitoes. PLOS Neglected Tropical Diseases, 11, e0005933.

Faulde, M.K., Spiesberger, M. & Abbas, B. (2012) Sentinel site-enhanced near-real time surveillance documenting West Nile virus circulation in two Culex mosquito species indicating different transmission characteristics, Djibouti City, Djibouti. Journal of the Egyptian Society of Parasitology, 42, 461–474.

Faye, O., Ba, Y., Faye, O., Talla, C., Diallo, D., Chen, R., et al. (2014) Urban Epidemic of Dengue Virus Serotype 3 Infection, Senegal, 2009 -Volume 20, Number 3—March 2014 - Emerging Infectious Diseases journal - CDC. Emerging Infectious Diseases, 20, 456–459.

Fenollar, F. & Mediannikov, O. (2018) Emerging infectious diseases in Africa in the 21st century. New Microbes and New Infections, 26, S10.

Florio, J., Verú, L.M., Dao, A., Yaro, A.S., Diallo, M., Sanogo, Z.L., et al. (2020) Diversity, dynamics, direction, and magnitude of high-altitude migrating insects in the Sahel. Scientific Reports 2020 10:1, 10, 1–14.

Foley, D.H., Rueda, L.M. & Wilkerson, R.C. (2007) Insight into Global Mosquito Biogeography from Country Species Records. Journal of Medical Entomology, 44, 554–567.

Fontenille, D., Traore-Lamizana, M., Diallo, M., Thonnon, J., Digoutte, J.P. & Zeller, H.G. (1998) New vectors of Rift Valley fever in West Africa. Emerging infectious diseases, 4, 289–293.

Garrett-Jones, C. (1950) A dispersion of mosquitoes by wind. Nature, 165, 285–285.

Garrett-Jones, C. (1962) The possibility of active long-distance migrations by Anopheles pharoensis Theobald. Bulletin of the World Health Organization, 27, 299–302.

Google Developers. (2021) Countries: Public data [WWW Document]. URL https://developers.google.com/public-data/docs/canonical/countries_csv [accessed on.

Guernier, V., Hochberg, M.E. & Guégan, J.F. (2004) Ecology drives the worldwide distribution of human diseases. PLoS Biology, 2.

Haddow, A.D., Nasar, F., Guzman, H., Ponlawat, A., Jarman, R.G., Tesh, R.B., et al. (2016) Genetic Characterization of Spondweni and Zika Viruses and Susceptibility of Geographically Distinct Strains of Aedes aegypti, Aedes albopictus and Culex quinquefasciatus (Diptera: Culicidae) to Spondweni Virus. PLOS Neglected Tropical Diseases, 10, e0005083.

Hanafi, H., Warigia, M., Breiman, R.F., Godsey, M., Hoel, D., Lutomiah, J., et al. (2010) Rift Valley Fever Virus Epidemic in Kenya, 2006/2007: The Entomologic Investigations. The American Journal of Tropical Medicine and Hygiene, 83, 28–37.

Hellgren, O., Waldenstrom, J., PerezERÉZ-Tris, J., Szoll, E., Si, Ö., Hasslequist, D., et al. (2007) Detecting shifts of transmission areas in avian blood parasites — a phylogenetic approach. Molecular Ecology, 16, 1281–1290.

Huestis, D.L., Dao, A., Diallo, M., Sanogo, Z.L., Samake, D., Yaro, A.S., et al. (2019) Windborne long-distance migration of malaria mosquitoes in the Sahel. Nature, 574, 404–408.

Irish, S.R., Kyalo, D., Snow, R.W. & Coetzee, M. (2020) Updated list of Anopheles species (Diptera: Culicidae) by country in the Afrotropical Region and associated islands. Zootaxa, 4747, 401–449.

Jones, K.E., Patel, N.G., Levy, M.A., Storeygard, A., Balk, D., Gittleman, J.L., et al. (2008) Global trends in emerging infectious diseases. Nature, 451, 990.

Joy, D.A., Gonzalez-Ceron, L., Carlton, J.M., Gueye, A., Fay, M., McCutchan, T.F., et al. (2008) Local adaptation and vector-mediated population structure in Plasmodium vivax malaria. Molecular Biology and Evolution, 25, 1245–1252.

Jupp, P. & McIntosh, B. (1990) Aedes furcifer and other mosquitoes as vectors of chikungunya virus at Mica, northeastern Transvaal, South Africa. J Am Mosq Control Assoc., 6, 415–420.

Jupp, P.G. (1996) Mosquitoes of Southern Africa: Culicinae and Toxorhynchitinae. Ekogilde Publishers.

Karabatsos, N. (1985) International Catalogue of Arboviruses, including certain other viruses of vertabrates. 3rd edn. Am Soc Trop Med Hygi for the Subcommittee on Information Exchange of the Am Comm on Arthro born viruses., San Antonio, Texas.

Kay, B.H. & Farrow, R.A. (2000) Mosquito (Diptera: Culicidae) Dispersal: Implications for the Epidemiology of Japanese and Murray Valley Encephalitis Viruses in Australia. Journal of Medical Entomology, 37, 797–801.

Kligler, I.J., Muckenfuss, R.S. & Rivers, T.M. (1928) Transmission of fowl-pox by mosquitoes. Proc. Soc. Exptl. Biol. Med., 26, 128–9.

Kyalo, D., Amratia, P., Mundia, C.W., Mbogo, C.M., Coetzee, M. & Snow, R.W. (2017) A geo-coded inventory of anophelines in the Afrotropical Region south of the Sahara: 1898-2016. Welcome Open Research, 2, 57-.

Laurence, B.R. (1989) The global dispersal of bancroftian filariasis. Parasitology Today, 5, 260– 264.

Lemine, A., Lemrabott, M., Ebou, M., Khadijetou, L., Salem, O., Khyarhoum, O., et al. (2017) Mosquitoes (Diptera: Culicidae) in Mauritania: A review of their biodiversity, distribution and medical importance. Parasites and Vectors, 10.

Lepage, D. (2021) Avibase - The World Bird Database [WWW Document]. URL https://avibase.bsc-eoc.org/checklist.jsp?region=AFC [accessed on.

Linder, H.P., Klerk, H.M. de, Born, J., Burgess, N.D., Fjeldså, J. & Rahbek, C. (2012) The partitioning of Africa: statistically defined biogeographical regions in sub-Saharan Africa. Journal of Biogeography, 39, 1189–1205.

Linthicum, K.J., Anyamba, A., Tucker, C.J., Kelley, P.W., Myers, M.F. & Peters, C.J. (1999) Climate and satellite indicators to forecast Rift Valley fever epidemics in Kenya. Science (New York, N.Y.), 285, 397–400.

Linthicum, K.J., Davies, F.G., Kairo, A. & Bailey, C.L. (1985) Rift Valley fever virus (family Bunyaviridae, genus Phlebovirus). Isolations from Diptera collected during an inter-epizootic period in Kenya. J Hyg (London), 95, 197–205.

Liu, W.M., Li, Y.Y., Learn, G.H., Rudicell, R.S., Robertson, J.D., Keele, B.F., et al. (2010) Origin of the human malaria parasite Plasmodium falciparum in gorillas. Nature, 467, 420–U67.

Lomolino, M. V. (2020) Biogeography A Very Short Introduction. First. Oxford University Press, Oxford, UK.

Molina-Cruz, A., Canepa, G.E., Kamath, N., Pavlovic, N. V., Mu, J., Ramphul, U.N., et al. (2015) Plasmodium evasion of mosquito immunity and global malaria transmission: The lock-and-key theory. Proceedings of the National Academy of Sciences, 112, 15178–15183.

Molina-Cruz, A., Canepa, G.E., Silva, T.L.A. e, Williams, A.E., Nagyal, S., Yenkoidiok-Douti, L., et al. (2020) Plasmodium falciparum evades immunity of anopheline mosquitoes by interacting with a Pfs47 midgut receptor. Proceedings of the National Academy of Sciences, 117, 2597– 2605.

Molina-Cruz, A., Garver, L.S., Alabaster, A., Bangiolo, L., Haile, A., Winikor, J., et al. (2013a) The human malaria parasite Pfs47 gene mediates evasion of the mosquito immune system. Science, 340, 984–987.

Molina-Cruz, A., Lehmann, T. & Knöckel, J. (2013b) Could culicine mosquitoes transmit human malaria? Trends in Parasitology.

Morse, S.S., Mazet, J.A., Woolhouse, M., Parrish, C.R., Carroll, D., Karesh, W.B., et al. (2012) Prediction and prevention of the next pandemic zoonosis. The Lancet, 380, 1956–1965.

Nanfack Minkeu, F. & Vernick, K.D. (2018) A Systematic Review of the Natural Virome of Anopheles Mosquitoes. Viruses 2018, Vol. 10, Page 222, 10, 222.

Ndiaye, E.H., Diallo, D., Fall, G., Ba, Y., Faye, O., Dia, I., et al. (2018) Arboviruses isolated from the Barkedji mosquito-based surveillance system, 2012-2013. BMC Infectious Diseases, 18, 1– 14.

Njabo, K.Y., Cornel, A.J., Sehgal, R.N.M., Loiseau, C., Buermann, W., Harrigan, R.J., et al. (2009) Coquillettidia (Culicidae, Diptera) mosquitoes are natural vectors of avian malaria in Africa. Malaria Journal, 8, 1–12.

Pedgley, D.E., Reynolds, D.R. & Tatchell, G.M. (1995) Long-range insect migration in relation to climate and weather: Africa and Europe. In Insect Migration: Tracking resources through space and time (ed. by Drake, V.A. & Gatehouse, A.G.). Cambridge University Press, New York, pp. 3–30.

Perkins, S.L. (2014) Malaria’s Many Mates: Past, Present, and Future of the Systematics of the Order Haemosporida. Journal of Parasitology, 100, 11–25.

Perkins, S.L. (2018) Malaria in Farmed Ungulates: an Exciting New System for Comparative Parasitology. mSphere, 3.

Phipps, W.L., López-López, P., Buechley, E.R., Oppel, S., Álvarez, E., Arkumarev, V., et al. (2019) Spatial and Temporal Variability in Migration of a Soaring Raptor Across Three Continents. Frontiers in Ecology and Evolution, 7.

Purdon, A., Mole, M.A., Chase, M.J. & Aarde R.J. van. (2018) Partial migration in savanna elephant populations distributed across southern Africa. Scientific Reports 2018 8:1, 8, 1–11.

R Core Team. (2016) R: A Language and Environment for Statistical Computing.

Reynolds, D.R., Chapman, J.W. & Harrington, R. (2006) The migration of insect vectors of plant and animal viruses. Advances in virus research, 67, 453–517.

Reynolds, D.R. & Riley, J.R. (1988) A migration of grasshoppers, particularly Diabolocatantops axillaris (Thunberg) (Orthoptera: Acrididae), in the West African Sahel. Bulletin of Entomological Research, 78, 251–271.

Ricklefs, R.E., Medeiros, M., Ellis, V.A., Svensson-Coelho, M., Blake, J.G., Loiselle, B.A., et al. (2017) Avian migration and the distribution of malaria parasites in New World passerine birds. Journal of Biogeography, 44, 1113–1123.

Rosenberg, R. (2015) Detecting the emergence of novel, zoonotic viruses pathogenic to humans. Cellular and Molecular Life Sciences, 72, 1115–1125.

Rosenberg, R., Johansson, M.A., Powers, A.M. & Miller, B.R. (2013) Search strategy has influenced the discovery rate of human viruses. Proceedings of the National Academy of Sciences of the United States of America, 110, 13961–13964.

Rutledge, G.G., Böhme, U., Sanders, M., Reid, A.J., Cotton, J.A., Maiga-Ascofare, O., et al. (2017) Plasmodium malariae and P. ovale genomes provide insights into malaria parasite evolution. Nature 2017 542:7639, 542, 101–104.

Sanogo, Z.L., Yaro, A.S., Dao, A., Diallo, M., Yossi, O., Samaké, D., et al. (2021) The effects of high-altitude windborne migration on survival, oviposition, and blood-feeding of the African malaria mosquito, Anopheles gambiae s.l. (Diptera: Culicidae). Journal of Medical Entomology, 58.

SAS Institute. (2012) SAS software for Windows Version 9.4.

Sellers, R.F. (1980) Weather, host and vector--their interplay in the spread of insect-borne animal virus diseases. The Journal of hygiene, 85, 65–102.

Service, M.W. (Ed.). (2001) Encyclopedia of Arthropod-transmitted Infections. 1st edn. CAB International., New York.

Seufi, A.E.M. & Galal, F.H. (2010) Role of Culex and Anopheles mosquito species as potential vectors of rift valley fever virus in Sudan outbreak, 2007. BMC Infectious Diseases, 10, 1–8.

Small, S.T., Labbé, F., Coulibaly, Y.I., Nutman, T.B., King, C.L., Serre, D., et al. (2019) Human Migration and the Spread of the Nematode Parasite Wuchereria bancrofti. Molecular Biology and Evolution, 36, 1931–1941.

Southwood, T.R.E. (1962) Migration of terrestrial arthropods in relation to habitat. Biological Reviews, 37, 171–211.

Swei, A., Couper, L.I., Coffey, L.L., Kapan, D. & Bennett, S. (2020) Patterns, Drivers, and Challenges of Vector-Borne Disease Emergence. Vector-Borne and Zoonotic Diseases, 20, 159–170.

Tandina, F., Doumbo, O., Yaro, A.S., Traoré, S.F., Parola, P. & Robert, V. (2018) Mosquitoes (Diptera: Culicidae) and mosquito-borne diseases in Mali, West Africa. Parasites & Vectors, 11, 467.

Tantely, L.M., Boyer, S. & Fontenille, D. (2015) A Review of Mosquitoes Associated with Rift Valley Fever Virus in Madagascar. The American Journal of Tropical Medicine and Hygiene, 92, 722.

Taylor, L.H., Latham, S.M. & Woolhouse, M.E.J. (2001) Risk factors for human disease emergence. Philosophical Transactions of the Royal Society B: Biological Sciences, 356, 983– 989.

The-World-Bank. (2021) Data: Land Area [WWW Document]. URL https://data.worldbank.org/indicator/AG.LND.TOTL.K2 [accessed on.

Tolley, K.A., Alexander, G.J., Branch, W.R., Bowles, P. & Maritz, B. (2016) Conservation status and threats for African reptiles.

Trari, B., Dakki, M. & Harbach, R.E. (2017) An updated checklist of the Culicidae (Diptera) of Morocco, with notes on species of historical and current medical importance. Journal of Vector Ecology, 42, 94–104.

Turell, M.J. & Knudson, G.B. (1987) Mechanical transmission of Bacillus anthracis by stable flies (Stomoxys calcitrans) and mosquitoes (Aedes aegypti and Aedes taeniorhynchus). Infection and Immunity, 55, 1859–1861.

Twohig, K.A., Pfeffer, D.A., Baird, J.K., Price, R.N., Zimmerman, P.A., Hay, S.I., et al. (2019) Growing evidence of Plasmodium vivax across malaria-endemic Africa. PLOS Neglected Tropical Diseases, 13, e0007140.

Vasilakis, N., Deardorff, E.R., Kenney, J.L., Rossi, S.L., Hanley, K.A. & Weaver, S.C. (2009) Mosquitoes put the brake on arbovirus evolution: Experimental evolution reveals slower mutation accumulation in mosquito than vertebrate cells. PLoS Pathogens, 5.

Vasilakis, N., Tesh, R.B., Popov, V.L., Widen, S.G., Wood, T.G., Forrester, N.L., et al. (2019) Exploiting the Legacy of the Arbovirus Hunters. Viruses 2019, Vol. 11, Page 471, 11, 471.

Veech, J.A. (2013) A probabilistic model for analysing species co-occurrence. Global Ecology and Biogeography, 22, 252–260.

Villinger, J., Mbaya, M.K., Ouso, D., Kipanga, P.N., Lutomiah, J. & Masiga, D.K. (2017) Arbovirus and insect-specific virus discovery in Kenya by novel six genera multiplex high-resolution melting analysis. Molecular Ecology Resources, 17, 466–480.

Wardeh, M., Risley, C., McIntyre, M.K., Setzkorn, C. & Baylis, M. (2015) Database of host-pathogen and related species interactions, and their global distribution. Scientific Data 2015 2:1, 2, 1–11.

Weaver, S.C., Chen, R. & Diallo, M. (2020) Chikungunya virus: Role of vectors in emergence from enzootic cycles. Annual Review of Entomology, 65, 313–332.

Weaver, S.C., Winegar, R., Manger, I.D. & Forrester, N.L. (2012) Alphaviruses: Population genetics and determinants of emergence. Antiviral Research, 94, 242–257.

WHO, W.H.O. (2020) Vector-borne diseases [WWW Document]. URL https://www.who.int/news-room/fact-sheets/detail/vector-borne-diseases [accessed on 2020].

Wilkerson, R.C., Linton, Y.-M. & Strickman, D. (2021) Mosquitoes of the World. Vol. 1 & 2. Johns Hopkins University Press, Baltimore.

Wolfe, N.D.N., Dunavan, C.C.P. & Diamond, J. (2007) Origins of major human infectious diseases. Nature, 447, 279–283.

Woolhouse, M.E.J. & Gowtage-Sequeria, S. (2005) Host range and emerging and reemerging pathogens. Emerging Infectious Diseases, 11, 1842–1847.

